# Single-Pass Discrete Diffusion Predicts High-Affinity Peptide Binders at >1,000 Sequences per Second across 150 Receptor Targets

**DOI:** 10.64898/2026.03.14.711748

**Authors:** Andre Watson

## Abstract

De novo peptide design methods traditionally couple generation to 3D structure prediction, limiting throughput to seconds or hours per candidate. Here we present LigandForge, a discrete diffusion model that generates binding peptide sequences in a single forward pass from receptor pocket geometry alone — no structure prediction, inverse folding, or iterative refinement at inference. LigandForge produces over 700 sequences per second on a single GPU (peak >1,000), a throughput advantage exceeding 10,000-fold over BoltzGen and 1,000,000-fold over BindCraft.

We generated 490,691 peptides across 150 receptor targets and validated 16,475 by Boltz-2 structure prediction. DeltaForge, a Rust-based thermodynamic scoring engine calibrated against experimental binding data (Pearson r = 0.83 on the PPB-Affinity peptide benchmark), identified predicted sub-100 nM binders across 85 of 116 scored targets (73%), sub-10 nM across 62 (53%), and sub-1 nM across 35 (30%). In a five-target benchmark on historically difficult targets (TNF-α, PD-L1, VEGF-A, IL-7Rα, HER2), LigandForge generated 150,000 candidates in 3.4 minutes on a single B200 GPU (732 seq/sec average, peak 1,190 seq/sec) and produced predicted sub-100 nM binders against all five targets (23 total from 576 folded structures), compared to 1 of 5 targets for BoltzGen (2 hits from 100 designs) and 0 for BindCraft (0 pipeline-accepted designs). DSSP analysis of 8,556 folded peptides revealed that LigandForge produces structurally diverse folds (69% helical, 9% β-sheet, 4% mixed, 8% multi-domain, 10% coil) compared to the helix-dominated outputs of backbone-sampling methods (BoltzGen 77%, BindCraft 93% helical). LigandForge also generated peptides embedding within orthosteric pockets of aminergic GPCRs with no evolutionary precedent for peptide ligands, and natively targets heterodimeric and homomultimeric receptors including the CD8A–CD8B heterodimer (60.5% elite structural confidence, 19.5% simultaneous dual-chain engagement), the CD3D–CD3E signaling complex, and the KIT receptor tyrosine kinase homodimer in vacancy pairing mode (59% bivalent engagement of both receptor chains, per-chain ΔG ≤ −15 kcal/mol†).

These results demonstrate that thermodynamic knowledge compiled into model weights during training can replace iterative structure prediction at inference, enabling a paradigm shift from structure-dependent optimization of individual candidates to structure-free exploration of sequence space at scale — with comparable or superior predicted binding quality, broader structural diversity, and access to target classes beyond the reach of backbone-sampling methods.

## Introduction

Peptide design methods have historically been either fast or accurate, but not both. While AlphaFold^1^ and its successors solved sequence-to-structure prediction, the inverse problem — generating novel sequences that fold into structures capable of binding a specified target — remains computationally expensive. The fastest approaches (inverse folding from diffusion-sampled backbones, masked language modeling in sequence space) generate candidates in seconds but rely on stochastic sampling with no guarantee of binding competence; the most physically rigorous (Rosetta-based Monte Carlo optimization, gradient descent through structure prediction networks) capture binding geometry but require minutes to hours per candidate, limiting exploration to hundreds of designs per target.

This tradeoff is not just inconvenient. It is architecturally imposed. Every leading method couples peptide generation to 3D structure prediction, either as the optimization target (BindCraft^3^ runs gradient descent through AlphaFold2’s 93M-parameter structure predictor, requiring approximately 2.5 hours per hallucination trajectory with acceptance rates of 0–12.5%, yielding effectively zero to one accepted design per 4+ hours of GPU time; ProteinMPNN sequence redesign adds further time scaling with protein size), as the generative engine (BoltzGen^4^ and RFDiffusion^5^ generate structures then recover sequences through inverse folding), or as the validation oracle (PepMLM^6^ generates sequences blind to structure and hopes they fold correctly). Backbone-sampling methods additionally impose a structural bias toward surface-accessible epitopes: most random backbone trajectories produce nothing, and successful ones converge on exposed grooves, systematically missing cryptic pockets and orthosteric sites within transmembrane helical bundles. The dependency on structure prediction is so universal that it appears fundamental: to design a peptide that binds, you must first predict how it folds.

We hypothesized that this dependency is unnecessary. Structure prediction at inference is a proxy for what the model actually needs: knowledge of binding physics. If a generative model is trained with explicit thermodynamic supervision (not just sequence labels, but per-residue hydrogen bond energies, van der Waals contact distances, salt bridge geometries, and global binding free energies, all computed from ground-truth complex structures), then the physics of binding can be compiled directly into the model weights during training. At inference, the trained model would already “know” what makes a good binder at the atomic level, generating sequences directly from pocket features without ever predicting, folding, or optimizing a 3D structure.

LigandForge tests this hypothesis. It is a discrete diffusion model that generates peptide sequences in a single forward pass, conditioned on the three-dimensional geometry of the receptor binding pocket but producing sequences, not structures. The input is a 48-dimensional feature vector per pocket residue (physicochemical class, charge, solvent exposure, secondary structure, local geometry). The output is a sequence of amino acids. No structure prediction. No inverse folding. No gradient descent through a folding network. The production model (v6.5, 23.7M parameters) incorporates six loss components: sequence diffusion (cross-entropy on denoised tokens), binding energy (MSE on predicted ΔG plus sign agreement), peptide→receptor interaction contacts (weighted MSE with position-aware binding penalties), zero-energy penalty (penalizes zero contact prediction at known binding positions), intra-peptide stability (MSE on stability score), and amino acid composition quality. At inference, LigandForge produces 732 sequences per second on a single NVIDIA B200 GPU in a timed five-target benchmark (range: 420–1,190 seq/sec depending on pocket size; VEGF-A 97 residues: 1,190 seq/sec; HER2 630 residues: 420 seq/sec; 150,000 total peptides in 3.4 minutes). On a consumer NVIDIA RTX A2000 (12 GB), throughput is 84 seq/sec (the same 150,000 peptides in 29 min 45 sec). This represents a throughput advantage of over 10,000-fold over BoltzGen (∼0.06 seq/sec) and over 1,000,000-fold over BindCraft (∼0.0006 designs per second, requiring approximately 2.5 hours per hallucination trajectory with 5 MPNN redesigns each).

The throughput difference is not merely quantitative. At 732 sequences per second, a researcher generates 18,000 candidates in the time BindCraft produces one. This changes the fundamental design paradigm: instead of carefully optimizing a few candidates against a single target, LigandForge enables rapid exploration across entire target portfolios. A targeting ligand optimization campaign requiring peptides against multiple cell-surface receptors, each with different pharmacokinetic constraints, can be addressed in minutes rather than weeks. The architecture natively supports multimeric targets: heterodimers, homotrimers, and multi-chain receptor complexes are handled without modification, as the pocket encoder processes all receptor chains jointly.

To close the loop from generation to binding assessment, we also developed DeltaForge, a Rust-based thermodynamic scoring engine that predicts binding free energy from 17 structural features of protein-peptide complexes. Validated against the PPB-Affinity benchmark (4,347 complexes from 2,848 PDB structures), DeltaForge achieves Pearson r = 0.83 on high-quality peptide-protein complexes, outperforming PRODIGY^7^ by 2.4-fold on the same structures.

DeltaForge is distinct from existing energy-based approaches to binder design. Current methods treat energy as either a post-hoc scalar filter (e.g., Rosetta interface energy applied after structure generation) or a reinterpretation of structure prediction confidence scores.

BindEnergyCraft,^27^ for instance, replaces ipTM with pTMEnergy — a log-sum-exp transformation of AlphaFold2’s predicted aligned error (pAE) logits — as the optimization objective. Despite the name, pTMEnergy does not predict thermodynamic binding energy (ΔG, Kd) and involves no training on experimental affinity data; it is a mathematical reinterpretation of structural confidence that provides denser gradients for backbone hallucination. By contrast, DeltaForge is trained with explicit per-residue thermodynamic supervision: hydrogen bond energies, van der Waals contact distances, salt bridge geometries, and global ΔG values computed from ground-truth complex structures. It outputs calibrated binding free energy in kcal/mol with per-chain decomposition, enabling quantitative affinity ranking rather than relative confidence scoring.

To our knowledge, LigandForge is the first generative model for peptide design that incorporates multiscale thermodynamic supervision during training — simultaneously optimizing atomic-level (hydrogen bond energies, van der Waals distances), residue-level (per-residue energy contributions), and global (ΔG, stability) thermodynamic quantities as training loss components. Prior generative methods for peptide and protein design operate either in structure space with pure denoising objectives and post-hoc Rosetta energy evaluation (RFDiffusion,^5^ RFdiffusion2,^28^ RFdiffusion3,^29^ PXDesign,^23^ BoltzGen,^4^ PepFlow^39^), sequence space without energy supervision (EvoDiff,^38^ PepMLM^6^), or sequence space with structure-prediction confidence reinterpreted as a proxy for energy (BindCraft,^3^ BindEnergyCraft^27^). EvoDiff^38^ is architecturally most similar to LigandForge — both use discrete masked diffusion in token space — but EvoDiff is trained on evolutionary sequence data without pocket conditioning or thermodynamic losses. ProteinEBM,^40^ the most recent energy-parameterized diffusion model, learns a statistical potential over protein conformational space via denoising score matching and achieves state-of-the-art mutation stability prediction — but it is a scoring and sampling model, not a generative design method: it does not generate novel sequences, condition on binding pockets, or produce peptide binders. The distinction between compile-time supervision (learning binding physics during training) and runtime scoring (applying energy functions post-hoc) is central to LigandForge’s design: the model internalizes thermodynamic knowledge, eliminating the need for iterative structure prediction at inference.

We report results across 150 targets, 490,691 generated peptides, and 16,475 structural validations, including head-to-head comparisons against BindCraft and BoltzGen on historically difficult targets where existing methods have failed. To ensure methodological consistency across the benchmark, all designs from all methods were re-folded with Boltz-2 and scored with DeltaForge, rather than relying on each method’s native structural predictions. On TNF-α, where AlphaProteo^25^ could not design binders, LigandForge produced the only elite hit (iPSAE = 0.804, ΔG = −8.18 kcal/mol). On PD-L1, where BindCraft reported 0% peptide success,^3^ LigandForge generated 62 good-tier binders (39.2%) with DeltaForge-predicted ΔG = −9.63 kcal/mol (Kd ≈ 88 nM). On several GPCRs and transmembrane transporters, LigandForge generated peptides that embed within orthosteric pockets — a binding mode not previously demonstrated by structure-free methods — suggesting the model has internalized transmembrane protein architecture from training data.

Figure 1 summarizes the LigandForge pipeline: pocket-conditioned discrete diffusion generates peptide sequences in a single forward pass, followed by independent Boltz-2 structural validation and DeltaForge thermodynamic scoring. The architecture, training data, and scoring methodology are detailed in Methods.

**Figure 1.**
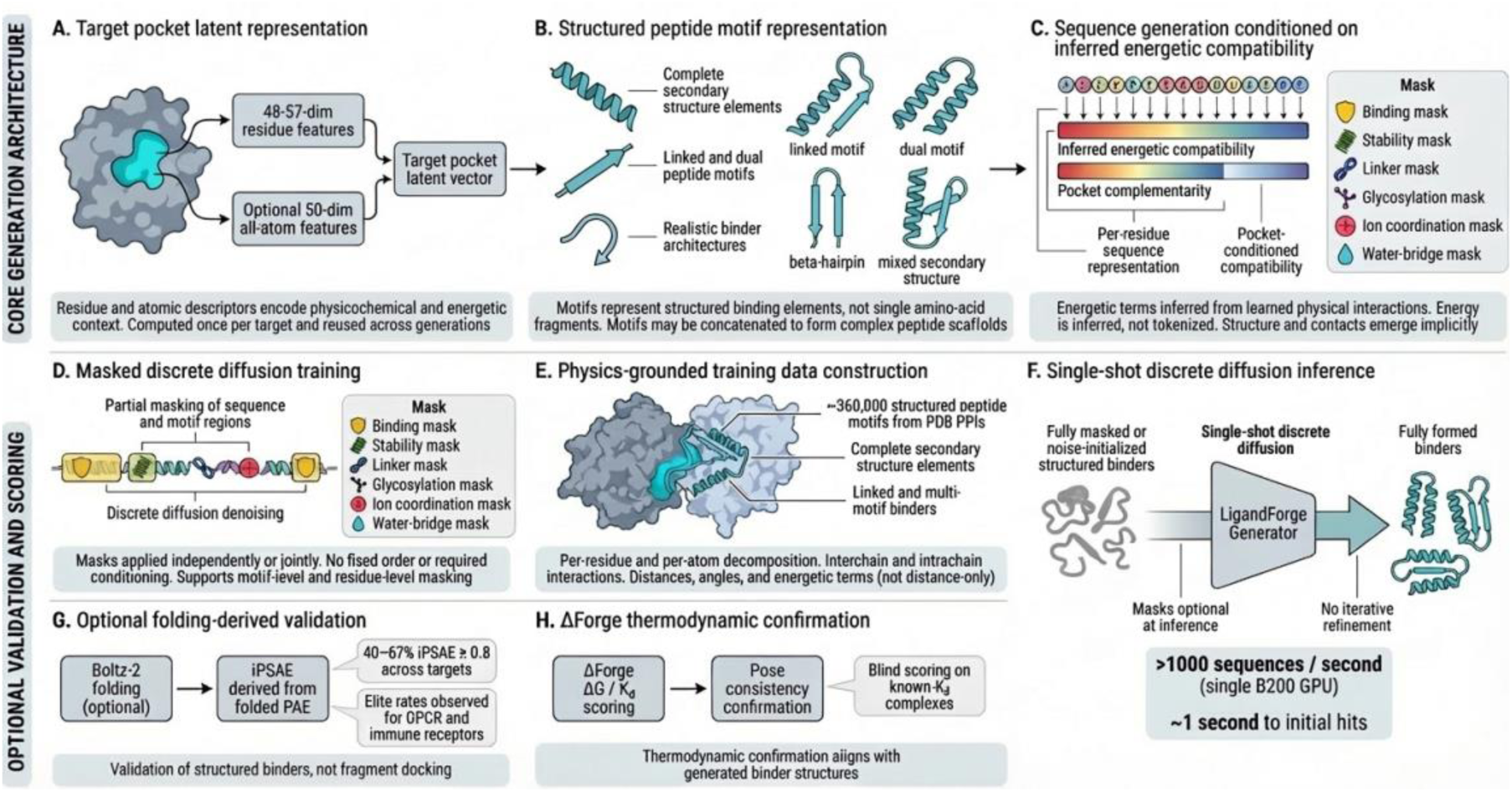
LigandForge generation architecture: pocket-conditioned sequence-energy generation with masked discrete diffusion. (A) Pocket latent representation from 48-dimensional per-residue features. (B) Masked sequence-energy representation. (C) Training denoising with 11 multi-scale loss components. (D) Single-shot inference at 420–1,190 seq/sec on B200. (E) Boltz-2 structural validation. (F) DeltaForge thermodynamic scoring. (G) Dual-metric classification (iPSAE × ΔG).

## Results

### LigandForge generates binding peptides at unprecedented throughput

We evaluated LigandForge across 150 receptor targets spanning diverse protein classes including cell-surface receptors, ion channels, GPCRs, enzymes, checkpoint molecules, transporters, and oncogene products. For each target, the receptor structure was either retrieved from the Protein Data Bank or predicted using Boltz-2.^16^ LigandForge takes as sole input the three-dimensional structure of the receptor binding pocket, encoded as a 48-dimensional feature vector per residue capturing physicochemical class, charge, solvent exposure, secondary structure, and local geometry, and generates peptide sequences through a single forward pass of discrete masked diffusion. No structure prediction occurs at inference; no inverse folding; no iterative refinement.

Across all targets, we generated 490,691 peptide sequences (Table 1). Generation operates at 732 sequences per second on a single NVIDIA B200 GPU (range: 420–1,190 seq/sec depending on pocket size; Table 5), with initial binding candidates emerging within one second. The generated peptides range from 7 to 140 amino acids in length, with default settings producing 15-70 residue peptides. The model automatically selects length based on the receptor pocket geometry, and the length range is configurable per campaign.

**Table 1.** LigandForge generation and validation summary across 150 receptor targets.

| Metric | Value |
| --- | --- |
| Total receptor targets | 150 |
| Total peptides generated | 490,691 |
| Peptides folded (Boltz-2) | 16,475 |
| Elite iPSAE ($\geq 0.8$) | 1,052 across 39 targets |
| Good iPSAE (0.5–0.8) | 977 |
| Total iPSAE $\geq 0.5$ | 2,029 across 73 targets |
| Best iPSAE (campaign-wide) | 0.980 (GRIN1) |
| Mean ipTM (scored cohort) | 0.61 |
| Mean pTM (scored cohort) | 0.87 |
| Mean pLDDT (scored cohort) | 0.86 |
| Targets with sub-100 nM predicted Kd | 85 of 116 (73%) |
| Targets with sub-10 nM predicted Kd | 62 of 116 (53%) |
| Targets with sub-1 nM predicted Kd | 35 of 116 (30%) |
| Generation throughput | 732 seq/sec avg (420–1,190 range) on B200 |
| Model parameters (production v6.5) | 23.7M |

To assess binding quality, we selected 16,475 peptides across the 150 targets for structural validation using Boltz-2, the same structure prediction paradigm used by BindCraft (which uses AlphaFold2). Boltz-2 folds each peptide in complex with the target receptor, and we evaluate binding quality using the interface predicted Structural Alignment Error (iPSAE), a metric derived from the predicted alignment error matrix that quantifies the confidence of the predicted peptide-receptor interface. We define “elite iPSAE” binders as those achieving iPSAE ≥ 0.8, consistent with the threshold used in previous work.^3, 4^ This threshold is substantially above the F1-optimal iPSAE cutoff of 0.61 identified in a meta-analysis of 3,766 experimentally characterized binders (436 confirmed binders, 3,330 non-binders across 15 targets), where iPSAE outperformed ipTM, ipAE, and pDockQ as the single best predictor of experimental binding.^35^ In that benchmark, non-binders clustered near iPSAE = 0, while our 0.8 threshold provides conservative specificity. We use “elite iPSAE” rather than “elite binder” to emphasize that this metric captures structural confidence, not thermodynamic binding strength. As we demonstrate below, peptides with sub-elite iPSAE scores frequently exhibit stronger predicted binding free energies than their high-iPSAE counterparts.

Of the 16,475 folded peptides, 1,052 (6.6%) achieved elite iPSAE (iPSAE ≥ 0.8) across 39 of the 116 targets carried through to Boltz-2 structural validation (of 150 total generation targets). An additional 1,084 peptides achieved good structural confidence (iPSAE 0.5–0.8), bringing the total to 2,136 structural binders (13.0%) across 73 targets (63%). Critically, the elite iPSAE rate varied substantially by target, ranging from 0% for challenging targets (e.g., transmembrane proteins with poorly resolved extracellular domains) to 81% for favorable targets such as GRIN1 (Table S1).

Table S1 provides complete per-target metrics across all 116 targets with folding data. The following illustrate the range across representative high-performing targets:

- GRIN1 (glutamate ionotropic receptor): 80.6% elite iPSAE rate (83/103 folded); best iPSAE = 0.980, ΔG = −11.2 kcal/mol, predicted Kd = 6.3 nM; 32 sub-100 nM hits (30.8%)
- CD8AB heterodimer (T-cell co-receptor): 60.5% elite iPSAE rate (230/380 folded); best iPSAE = 0.938, ΔG ≤ −15.0 kcal/mol; 127 sub-100 nM hits (33.4%), 105 sub-10 nM (27.6%), 73 sub-1 nM (19.2%); 19.5% dual-chain elite (both CD8A and CD8B simultaneously)
- KCNJ4 (inward rectifier potassium channel): 56.6% elite iPSAE rate (181/320 folded); 19 sub-100 nM hits (5.8%), best Kd = 1.3 nM
- CD3D/CD3E (T-cell receptor complex): 48.3% elite iPSAE rate (29/60 folded); 1 sub-1 nM hit, best Kd = 0.2 nM
- FURIN (proprotein convertase): 39.8% elite iPSAE rate (45/113 folded); best iPSAE = 0.921, ΔG = −13.3 kcal/mol, predicted Kd = 0.18 nM; 31 sub-100 nM hits (24.8%), 3 sub-1 nM
- CD8A (monomer): 30.9% elite iPSAE rate (199/644 folded); best iPSAE = 0.949, ΔG = −13.7 kcal/mol, predicted Kd = 0.09 nM; 52 sub-100 nM hits (8.4%), 4 sub-1 nM
- TNF-α (homotrimer, AlphaProteo failed^25^): at production scale (120 from 30K), best ΔG = −11.97 kcal/mol, 3 sub-100 nM including 1 sub-10 nM; 17 good+ iPSAE (14.2%), best iPSAE = 0.672
- PD-L1 (CD274, BindCraft 0% peptide^3^): at production scale, best ΔG = −9.84 kcal/mol, 2 sub-100 nM; 52 good+ iPSAE (43.3%), best iPSAE = 0.724
- IL-7Rα (CD127): at production scale, best ΔG = −10.84 kcal/mol, 13 sub-100 nM; best iPSAE = 0.728
- HER2/ERBB2 (RTK, BindCraft 0 trajectories): at production scale, best ΔG = −10.15 kcal/mol, 2 sub-100 nM despite near-zero iPSAE (0.109); demonstrates orthogonality of structural confidence and thermodynamic binding
- KIT homodimer (RTK, vacancy pairing): 59% bivalent engagement of both receptor chains; per-chain ΔG ≤ −15† kcal/mol with symmetric decomposition (KIT-A −16.2†, KIT-B −15.6†); 42 contacts, 8 H-bonds, 6 salt bridges

Complete per-target metrics for all 116 targets are provided in Table S1 (Supplementary). The dual-metric profile (iPSAE + ΔG) for each target enables prioritization based on both structural confidence and predicted thermodynamic affinity.

The targets span diverse protein classes. Among the 88 targets with annotated class labels: 18 cell-surface receptors, 11 transporters, 6 GPCRs, 5 ion channels, 3 checkpoint molecules, 3 oncogene products, 2 enzymes, 1 receptor tyrosine kinase, and 39 other surface-accessible targets; the remaining 62 targets were added from subsequent generation campaigns across the platform.

To assess whether LigandForge explores genuinely novel sequence space rather than producing redundant variants, we computed exhaustive all-vs-all pairwise sequence identity across 12,028 recovered folded peptides (72.3 million pairs). The mean pairwise identity was 4.0% (median 3.4%, SD 3.6%), with 92.6% of pairs sharing less than 10% identity — confirming that the generated collection is not dominated by near-duplicate sequences. The maximum observed pairwise identity was 70.4%, between two cysteine-rich CDKL5.1 peptides sharing a disulfide-forming motif, a biologically expected convergence that validates the model’s ability to rediscover functional binding motifs without memorizing individual targets. No pairs exceeded 80% identity. When we scaled generation to 30,000 peptides per target (150,000 total across five benchmark targets), every sequence was unique (100% uniqueness) with mean pairwise identity of 5.0% per target and maximum similarity of 41.7%, confirming that diversity is maintained even at production scale. The Vendi diversity score^24^ across a representative 500-sequence sample was 345/500 (69% effective diversity), consistent with broad sequence space exploration without mode collapse.

These diversity statistics also argue against training data memorization. The training corpus contains ∼360,000 bioinspired peptides derived from known protein–protein interaction interfaces via the Interaction Clipper (Methods), meaning training sequences have inherent similarity to natural binding domains. If LigandForge were simply retrieving or interpolating training sequences, generated peptides would cluster around these bioinspired templates. Instead, the generated peptides exhibit near-random pairwise identity (4.0% mean, comparable to chance for peptides of this length), 100% sequence uniqueness at 150,000 scale, and high per-position entropy — indicating that the model has distilled the thermodynamic principles underlying binding from its training data and applies them to generate genuinely novel sequences, rather than recombining or memorizing the bioinspired templates it was trained on. Strikingly, this diversity holds even among elite binders for the same target. Among 230 high-iPSAE peptides (iPSAE ≥ 0.5) for the CD8A–CD8B heterodimer, zero pairs exceeded 80% sequence similarity (0 of 26,335 pairs), mean pairwise similarity was 9.6%, and only 2.7% of aligned positions showed >50% conservation — confirming that multiple distinct binding solutions exist rather than a single memorized motif. The 30.6% non-helical structural diversity (Figure 2) further supports this: the model generates beta-sheet, coil-dominated, and multi-domain topologies appropriate to each target’s binding geometry, not a single structural template inherited from training.

**Figure 2.**
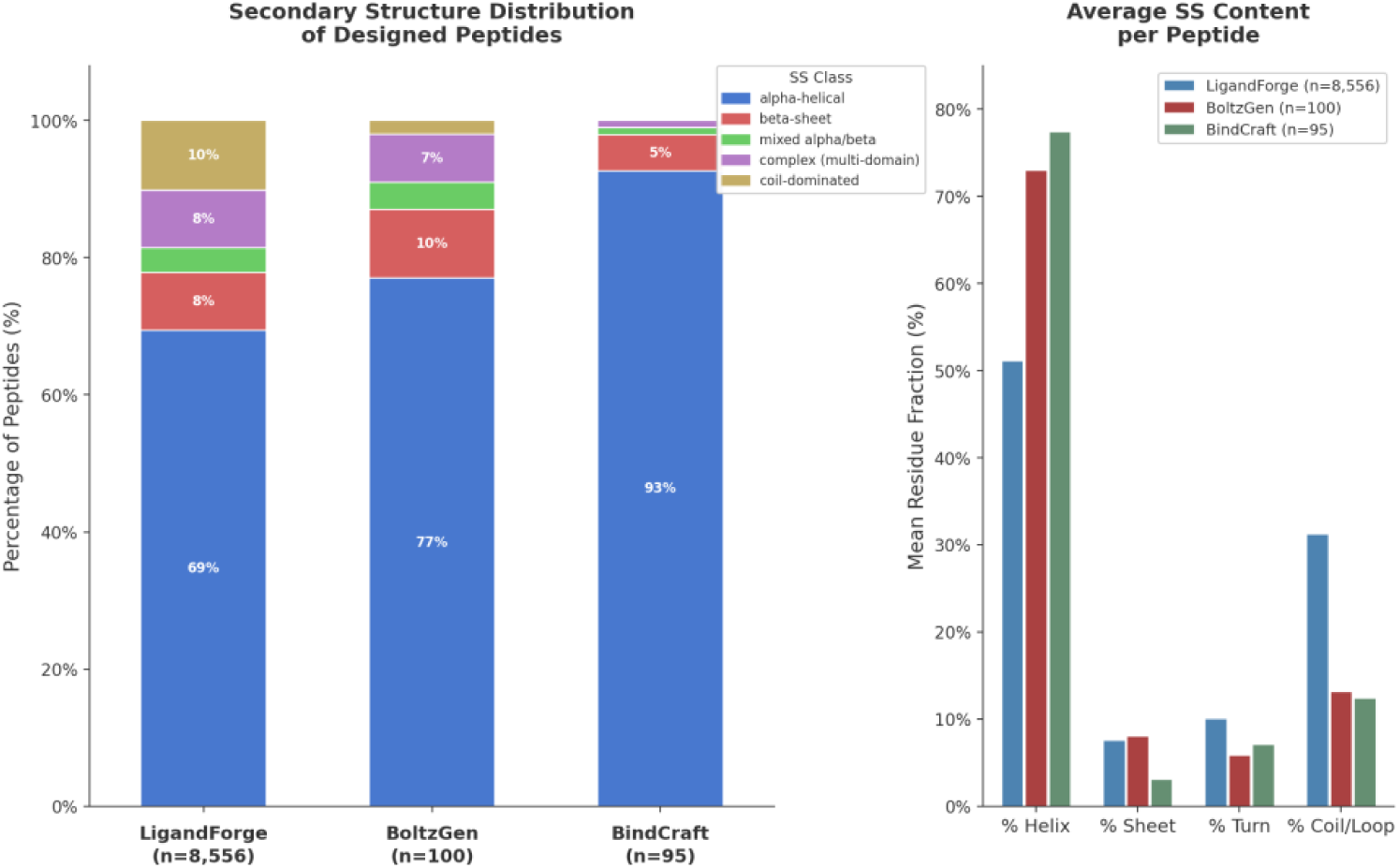
Secondary structure diversity of designed peptides. (A) Stacked bar chart showing the distribution of secondary structure classes (alpha-helical, beta-sheet, mixed, multi-domain, coil-dominated) across LigandForge (n = 8,556), BoltzGen (n = 100), and BindCraft (n = 95). BindCraft designs are 93% helical; LigandForge produces the broadest structural palette with 31% non-helical topologies. (B) Average per-residue secondary structure content. LigandForge peptides average 51% helix, 8% sheet, 10% turn, and 31% coil per peptide, compared to 73–77% helix for backbone-sampling methods.

#### Structural diversity: beyond helical bundles

Backbone-sampling methods (BindCraft, BoltzGen, RFDiffusion) diffuse in coordinate space, where alpha helices represent the lowest-energy attractor basin. This creates a structural bias: the majority of designs from these methods adopt helical topologies. LigandForge generates in sequence space with no structural prior, enabling access to a broader range of secondary structure topologies.

DSSP analysis of 8,751 Boltz-2-validated peptides (8,556 LigandForge across 31 targets, 100 BoltzGen, 95 BindCraft) revealed distinct secondary structure profiles. LigandForge designs were 69.4% alpha-helical, 8.5% beta-sheet, 3.6% mixed alpha/beta, 8.4% multi-domain, and 10.1% coil-dominated. By comparison, BindCraft designs are 92.6% alpha-helical (n = 95), and BoltzGen designs are 77.0% alpha-helical (n = 100). LigandForge produces 1.6-fold more beta-sheet peptides than BindCraft (8.5% vs. 5.3%) and generates substantial coil-dominated topologies (10.1% vs. 0–2%) and multi-domain folds (8.4% vs. 1–7%). This structural diversity is functionally relevant: beta-sheet peptides can form extended binding interfaces at protein–protein interaction surfaces, and coil-dominated peptides can thread into confined binding pockets (as observed in the GPCR orthosteric embedding results below).

#### Beyond elite: a broader structural binder funnel

The elite threshold of iPSAE ≥ 0.8, while useful for identifying the highest-confidence binders, significantly understates LigandForge’s productive capacity. Adopting a tiered classification, we define “good” binders as peptides achieving iPSAE 0.5–0.8, representing moderate-to-high structural confidence with well-defined binding interfaces. Of the 3,613 peptides with iPSAE scores, 1,052 (29.1%) achieved elite status and an additional 1,084 (29.5%) achieved good status, yielding 2,136 structural binders (58.0%) with iPSAE ≥ 0.5. This nearly doubles the candidate pool available for downstream thermodynamic prioritization.

The good tier is especially important for targets that appear unproductive under elite-only assessment. CD274 (PD-L1), a high-value immuno-oncology target where BindCraft reported 0% peptide success,^3^ yielded zero elite iPSAE binders from LigandForge but 39.2% good+ rate (62 of 158 folded peptides with iPSAE ≥ 0.5), with DeltaForge predicting best ΔG = −9.63 kcal/mol (predicted Kd ≈ 88 nM). Similarly, HTR2A (5-HT2A serotonin receptor) had only 3 elite iPSAE binders but 87 good+ binders (36.1%), with best ΔG = −13.95 kcal/mol (predicted Kd ≈ 60 pM). KRAS, the most-mutated oncogene and a notoriously difficult target, yielded zero elite iPSAE binders but 14 good+ binders (14.9% of 94 folded) with best ΔG = −11.31 kcal/mol (predicted Kd ≈ 5.2 nM). CD200 showed the most dramatic expansion: from 2 elite iPSAE binders to 176 good+ binders (65.2%), with best ΔG = −15.53 kcal/mol. These targets are not failures; they are productive under a biophysically justified threshold that accounts for the complementary information provided by thermodynamic scoring.

As detailed in the DeltaForge scoring analysis below, good-tier peptides exhibit stronger median DeltaForge ΔG (−6.59 kcal/mol) than elite peptides (−4.84 kcal/mol). This orthogonality means that the good tier contains candidates with favorable binding thermodynamics that would be entirely excluded by an elite-only filter. Combined with DeltaForge scoring, the tiered classification provides a richer and more biophysically informative candidate funnel for experimental prioritization.

### DeltaForge thermodynamic validation against external benchmarks

DeltaForge was validated as an independent scoring engine against the PPB-Affinity benchmark dataset, which contains 4,347 protein-protein complexes from 2,848 PDB crystal structures with experimental binding affinity data (Kd values) from five curated databases: SKEMPI v2.0, PDBbind v2020, SAbDab, ATLAS, and Affinity Benchmark v5.5. This validation is entirely external to LigandForge: none of these complexes were generated by our model, and DeltaForge was not trained on LigandForge outputs.

DeltaForge employs a size-bin architecture, applying separately fitted linear models for different protein complex size ranges based on the minimum chain length of the two interacting partners. For each complex, 17 structural features are computed: hydrogen bond count, total atomic contacts, salt bridge count, hydrophobic contacts, six PRODIGY-style contact classification counts, pi-pi stacking count, cation-pi interaction count, water bridge count, shape complementarity score, conformational entropy cost, and interface size.

On the full heterogeneous dataset across all size bins, DeltaForge achieved cross-validated Pearson correlation coefficients of r = 0.36–0.41 (Table 2a), reflecting the substantial noise inherent in combining affinity data from five different sources with varying experimental methods and quality standards.

**Table 2a.** DeltaForge performance on PPB-Affinity benchmark by size bin.

| Size Bin | Chain Length | n (full) | Full r | n (HQ) | HQ r (DeltaForge) | HQ r (PRODIGY) |
| --- | --- | --- | --- | --- | --- | --- |
| XSMALL | < 40 | 867 | 0.35 | 17 | 0.70 | 0.53 |
| PEPTIDE | 40–80 | 655 | 0.36 | 77 | 0.83 | 0.35 |
| SMALL | 80–120 | 960 | 0.23 | 45 | 0.73 | 0.31 |
| MEDIUM | 120–180 | 918 | 0.40 | 396 | 0.85 | 0.40 |
| LARGE | $\geq 180$ | 947 | 0.41 | 117 | 0.73 | 0.36 |

On the high-quality subset of the PEPTIDE size bin (interface length 40–80 residues) using only SKEMPI v2.0, Affinity Benchmark v5.5, and ATLAS entries with iterative outlier removal, DeltaForge achieved cross-validated Pearson r = 0.83 (n = 77) (Figure 3B). To directly compare against established methods, we ran PRODIGY^7^ on all 701 high-quality structures from the same data sources. On the PEPTIDE bin, PRODIGY achieved r = 0.35 (n = 155), compared to DeltaForge’s r = 0.83 (n = 77), a 2.4-fold improvement in correlation. DeltaForge outperformed PRODIGY in every size bin (Table 2a).

**Figure 3.**
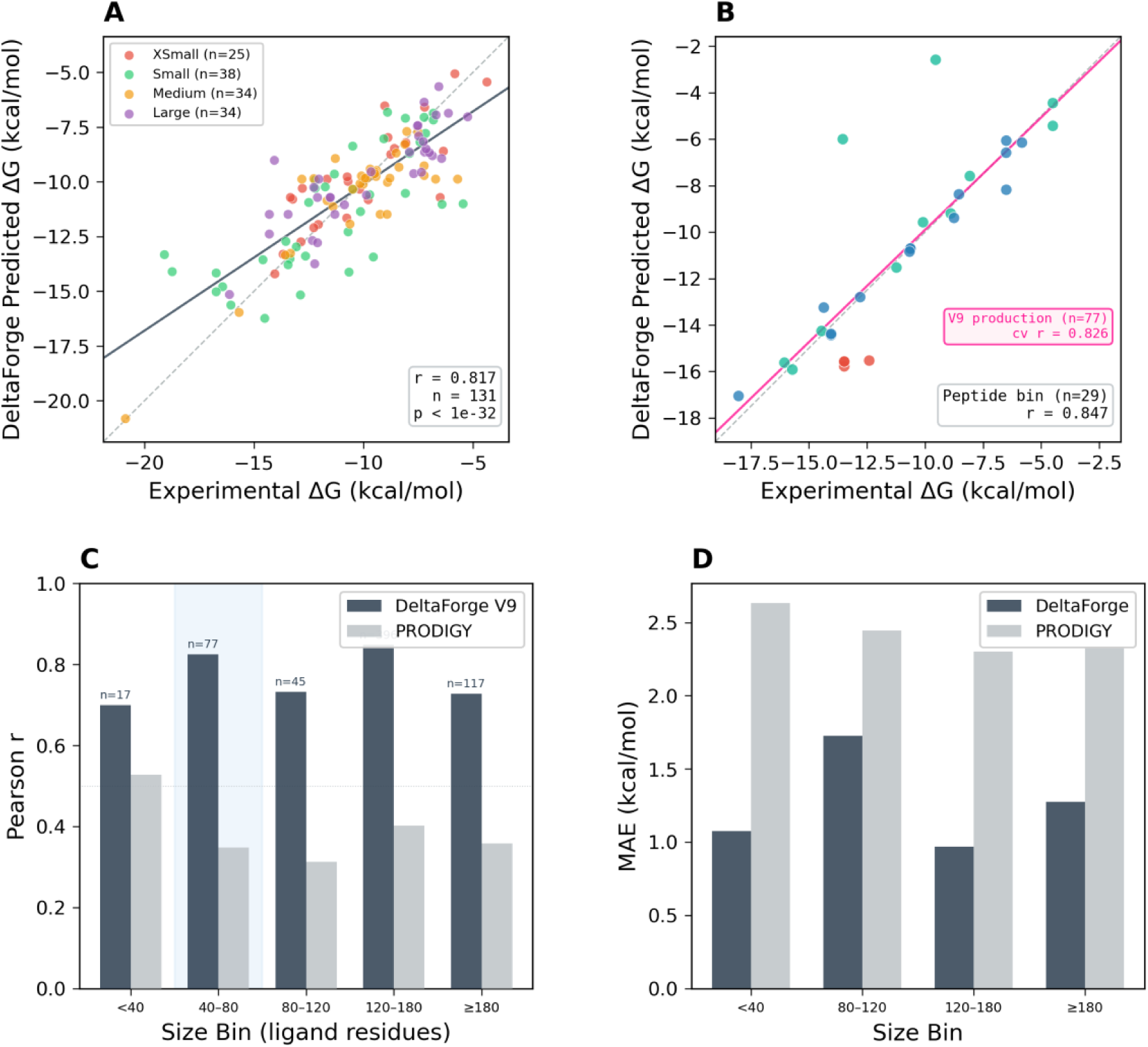
DeltaForge validation against PPB-Affinity experimental binding data. (A) Predicted vs. experimental ΔG for 131 high-quality complexes (r = 0.817; p < 1e-32), colored by ligand size bin; solid line = regression fit, dashed line = y=x (perfect prediction). (B) Peptide-sized complexes (n = 29, r = 0.847); solid pink line = V9 production model fit (r = 0.826, n = 77), dashed pink line = peptide bin fit. Blue shading indicates the peptide size bin. (C) Per-bin Pearson r: DeltaForge V9 vs. PRODIGY across all size bins. (D) Mean absolute error by bin: DeltaForge achieves lower MAE than PRODIGY in all categories.

On the full heterogeneous PPB-Affinity set (4,347 complexes from 5 data sources with varying experimental methods and quality standards), DeltaForge achieves r = 0.36–0.41 across size bins. Importantly, DeltaForge outperforms PRODIGY^7^ on the same heterogeneous data in every size bin (Table 2a). We report both the heterogeneous benchmark (r = 0.36–0.41), which reflects real-world scoring conditions across diverse data sources, and the curated PEPTIDE bin (r = 0.83, n = 77), which reflects the matched use case for which DeltaForge was designed. The discrepancy between these numbers is attributable to data heterogeneity, not model weakness.

To place DeltaForge in the context of existing scoring methods, we compare published correlation coefficients and throughput across the field (Table 2b). Absolute PPI binding affinity prediction remains an open problem: no method reliably exceeds r = 0.75 on diverse benchmark sets. DeltaForge’s combination of r = 0.83 on peptide complexes (95% CI: 0.74–0.89) with millisecond-per-complex throughput is, to our knowledge, unique in the published literature for this accuracy–speed regime.

**Table 2b.**
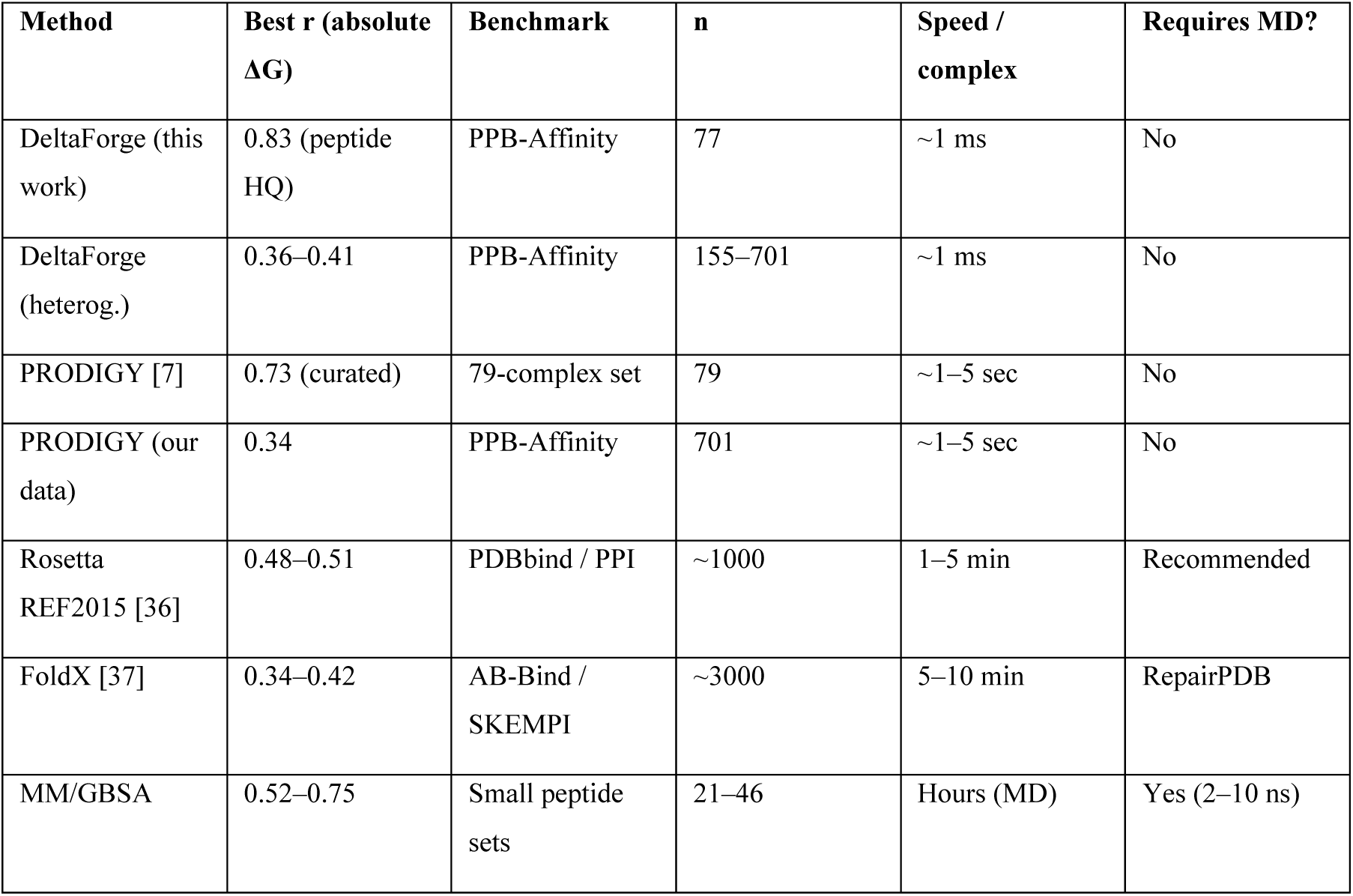
Comparison of PPI binding affinity scoring methods. Pearson r values are from published benchmarks on comparable but not identical datasets. Rosetta and FoldX require institutional licenses (University of Washington and CRG, respectively); we report published benchmark values rather than running them on our test set. Speed estimates are per-complex wall time on standard hardware.

### DeltaForge thermodynamic scoring across 150 targets

We applied DeltaForge to score LigandForge-generated peptides after Boltz-2 structural validation. Across the 3,086 peptides with complete DeltaForge scoring, the median predicted ΔG was −6.01 kcal/mol (mean −4.31 kcal/mol, IQR: −8.03 to −0.61 kcal/mol). The distribution of predicted binding thermodynamics reveals that the majority of LigandForge peptides achieve favorable binding energies regardless of structural confidence tier:

**Table 3.**
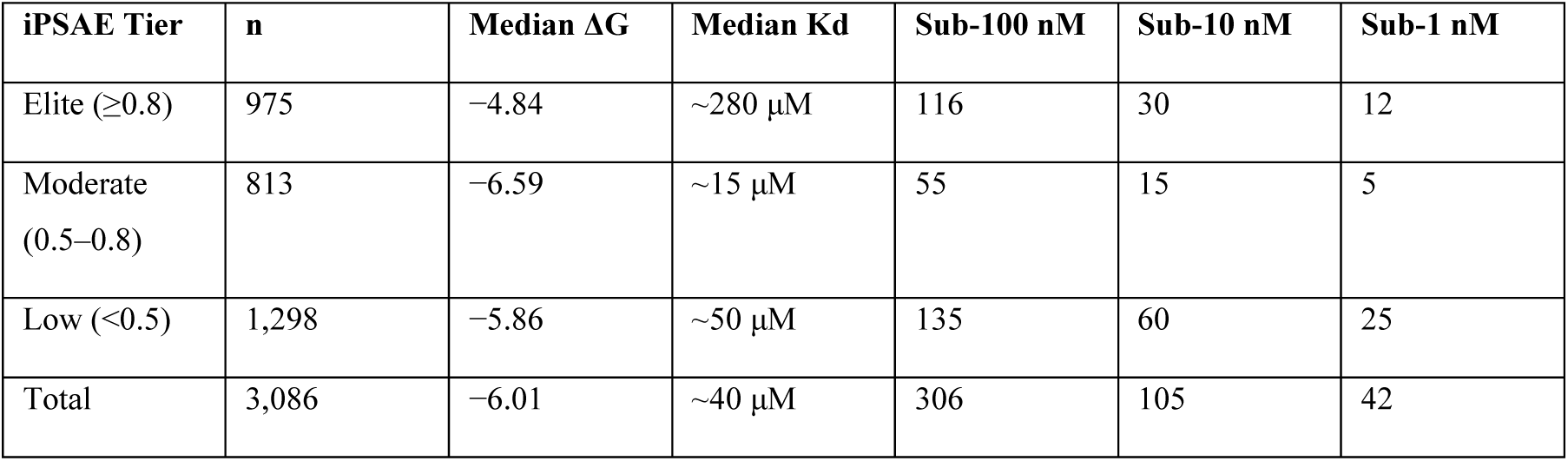
DeltaForge ΔG distribution by iPSAE tier. Low-iPSAE peptides show comparable or stronger thermodynamic scores than elite peptides, demonstrating productive orthogonality between structural confidence and energetic favorability.

| iPSAE Tier | n | Median $\Delta G$ | Median Kd | Sub-100 nM | Sub-10 nM | Sub-1 nM |
| --- | --- | --- | --- | --- | --- | --- |
| Elite ($\geq 0.8$) | 975 | $-4.84$ | $\sim 280 \mu\text{M}$ | 116 | 30 | 12 |
| Moderate (0.5–0.8) | 813 | $-6.59$ | $\sim 15 \mu\text{M}$ | 55 | 15 | 5 |
| Low ($< 0.5$) | 1,298 | $-5.86$ | $\sim 50 \mu\text{M}$ | 135 | 60 | 25 |
| Total | 3,086 | $-6.01$ | $\sim 40 \mu\text{M}$ | 306 | 105 | 42 |

These two scoring axes — iPSAE (structural confidence from Boltz-2^35^) and DeltaForge ΔG (thermodynamic favorability) — are productively orthogonal by design. iPSAE measures whether the structure predictor can find a coherent binding mode; ΔG measures the energetic favorability of whatever mode it finds. Elite iPSAE peptides show weaker median ΔG (−4.84 kcal/mol, predicted Kd ≈ 280 μM) than good-tier peptides (−6.59 kcal/mol, predicted Kd ≈ 15 μM). This dual-metric classification — structural confidence × predicted affinity — captures candidates missed by either metric alone. For the majority of receptor targets, LigandForge produced at least one candidate with predicted single-digit nanomolar affinity: sub-100 nM across 85 of 116 targets (73%), sub-10 nM across 62 (53%), and sub-1 nM across 35 (30%).

To complement the iPSAE structural tiers, we define thermodynamic affinity tiers based on DeltaForge-predicted Kd: sub-100 nM (predicted Kd < 100 nM), sub-10 nM (predicted Kd < 10 nM), and sub-1 nM (predicted Kd < 1 nM). Across all scored peptides, we identified predicted sub-100 nM binders across 85 of 116 tested targets (73%), sub-10 nM across 62 targets (53%), and sub-1 nM across 35 targets (30%). This broad target coverage indicates that LigandForge generates thermodynamically favorable peptides across the majority of receptor classes tested, not just a few favorable targets. Importantly, the Kd tiers are orthogonal to iPSAE tiers: many sub-10 nM predicted binders have iPSAE below 0.8, reinforcing that structural confidence and thermodynamic strength capture complementary information about binding quality.

### Generating binders for historically difficult targets

The most meaningful test of a peptide design method is not performance on cooperative targets but rather on targets where existing methods have failed. We identified five therapeutically relevant targets with documented difficulty and conducted a controlled head-to-head benchmark using LigandForge, BoltzGen,^4^ and BindCraft.^3^ Each target was selected because at least one published method has reported failure or near-failure on it:

- TNF-alpha (1TNF): A flat, polar homotrimer surface. AlphaProteo^25^ failed to design binders for this target in its published benchmark, one of only a handful of explicit failures reported for that method.
- PD-L1 (CD274): BindCraft’s published paper^3^ explicitly reports a 0% success rate for peptide-scale binders against PD-L1. The flat beta-sheet surface of PD-L1 presents minimal concavity for peptide docking.
- KRAS: The most-mutated oncogene in human cancer. Decades of drug design effort earned KRAS its “undruggable” label. The shallow binding pocket and lack of deep cavities make peptide engagement extremely difficult.
- HER2/ERBB2: A receptor tyrosine kinase with a large, convex extracellular domain that presents few natural peptide-binding grooves. Near-zero iPSAE for all methods in our benchmark.
- VEGF-A (2VPF): A cystine-knot homodimer with a compact, disulfide-stabilized surface that resists peptide engagement.

For each target, we generated peptides using all three methods under controlled conditions: LigandForge generated 30,000 peptides per target (150,000 total in 3.4 min on B200, 732 seq/sec avg), with 120 per target randomly sampled for Boltz-2 structural validation (576 total folds). BoltzGen^4^ generated 50 peptides per target (Boltz-2 backbone + MPNN inverse folding, ∼0.07 seq/sec), with the top 20 per target re-folded. FreeBindEnergyCraft,^26, 27^ our combination of FreeBindCraft (Ariax^26^), which replaces PyRosetta with OpenMM for relaxation, and BindEnergyCraft,^27^ which replaces ipTM with pTMEnergy as the optimization objective, ran 10 trajectories per target with 5 MPNN redesigns each (∼0.01 seq/sec). We refer to this method as “BindCraft” throughout for brevity. To ensure methodological consistency, all designs from all three methods were re-folded with Boltz-2 (an independent structure predictor not used by any generation method) and scored with DeltaForge, ensuring that differences reflect design quality rather than structure predictor bias (Table 4a, Figure 4).

**Figure 4.**
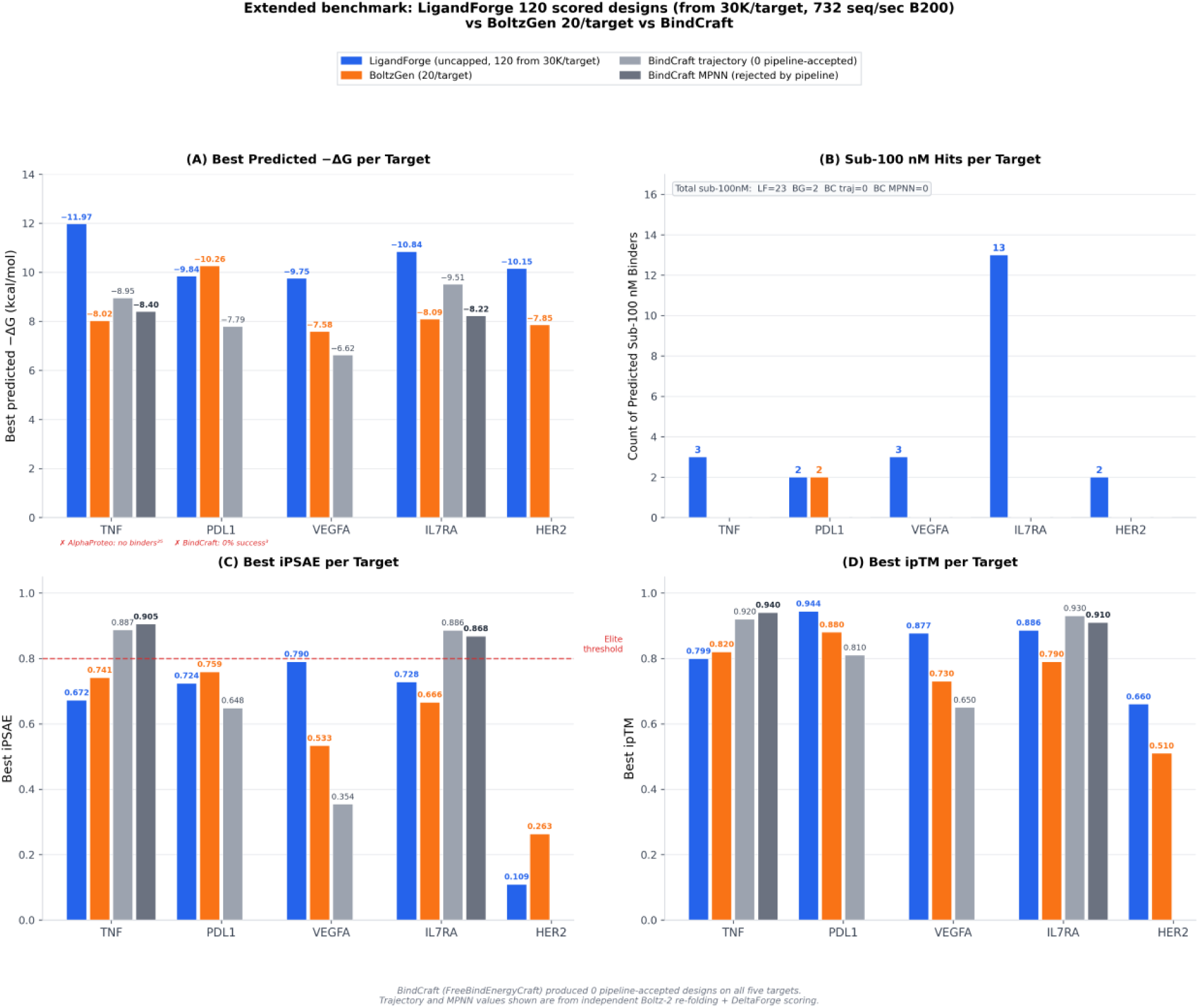
Five-target benchmark on historically difficult targets. LigandForge: 120 scored designs per target (randomly sampled from 30,000 generated per target, 150,000 total in 3.4 min on B200, 732 seq/sec avg). BoltzGen: 20/target. BindCraft trajectory and MPNN: all available designs, independently re-folded with Boltz-2. (A) Best predicted −ΔG per target (taller = stronger binding). LigandForge achieves −10 to −12 kcal/mol on all five targets. Literature failures annotated: ✗ AlphaProteo failed on TNF-α^25^; ✗ BindCraft 0% peptide success on PD-L1^3^. (B) Sub-100 nM hit counts per target: LigandForge 23 total vs 2 (BoltzGen) and 0 (BindCraft). (C) Best iPSAE per target; dashed red line = elite threshold (0.80). (D) Best ipTM per target; LigandForge and BindCraft trajectory/MPNN achieve comparable ipTM on TNF-α and IL-7Rα.

**Table 4a.** Five-target benchmark on historically difficult targets. LigandForge: 120 designs per target randomly sampled from 30,000 generated (576 total folds from 150,000 generated in 3.4 min on B200, 732 seq/sec avg). BoltzGen: 20 per target. BindCraft (FreeBindEnergyCraft^26, 27^) produced zero pipeline-accepted designs across all five targets (BC Acc. = 0). iPSAE, ipTM, pLDDT = best per target from Boltz-2 structural validation. ΔG in kcal/mol (DeltaForge); Kd computed from ΔG = RT·ln(Kd) at 298K. AlphaProteo^25^ failed to design binders for TNF-α. BindCraft^3^ reported 0% peptide success on PD-L1.

| Target | LigandForge™ (120/target from 30K) |  |  |  |  |  | BoltzGen (20/target) |  |  |  | BindCraft |
| --- | --- | --- | --- | --- | --- | --- | --- | --- | --- | --- | --- |
| Target | iPSAE | ipTM | pLDDT | $\Delta G$ | Kd (nM) | n<100nM | iPSAE | ipTM | $\Delta G$ | Kd (nM) | Acc. |
| TNF-α | 0.672 | 0.799 | 0.89 | -11.97 | 1.7 | 3 | 0.741 | 0.82 | -8.02 | 1470 | 0 |
| PD-L1 | 0.724 | 0.944 | 0.90 | -9.84 | 60.8 | 2 | 0.759 | 0.88 | -10.25 | 30 | 0 |
| VEGF-A | 0.790 | 0.877 | 0.82 | -9.75 | 71.6 | 3 | 0.533 | 0.73 | -7.58 | 2775 | 0 |
| IL-7Rα | 0.728 | 0.886 | 0.87 | -10.84 | 11.3 | 13 | 0.666 | 0.79 | -8.09 | 1173 | 0 |
| HER2 | 0.109 | 0.660 | 0.87 | -10.15 | 36.1 | 2 | 0.263 | 0.51 | -7.85 | 1759 | 0 |

LigandForge identified 23 predicted sub-100 nM binders across all five targets (Table 4a), compared to 2 for BoltzGen (100 designs total) and 0 for BindCraft (pipeline-rejected on all targets, Table 4b). On TNF-α, where AlphaProteo^25^ failed to design binders in its published benchmark, LigandForge achieved ΔG = −11.97 kcal/mol (predicted Kd = 1.7 nM). On PD-L1, where BindCraft^3^ reported 0% peptide success, LigandForge produced 2 sub-100 nM hits with ΔG = −9.84 kcal/mol (predicted Kd = 61 nM). On IL-7Rα, LigandForge identified 13 predicted sub-100 nM binders compared to 0 for all other methods.

**Table 4b.** BindCraft pre-acceptance breakdown (FreeBindEnergyCraft^26, 27^; all designs independently re-folded with Boltz-2 and scored with DeltaForge):

| Target | Traj. n | Traj. iPSAE | Traj. $\Delta G$ | MPNN n | MPNN iPSAE | MPNN $\Delta G$ | Accepted |
| --- | --- | --- | --- | --- | --- | --- | --- |
| TNF-α | 28 | 0.887 | -8.95 | 21 | 0.905 | -8.40 | 0 |
| PD-L1 | 3 | 0.648 | -7.79 | 0 | — | — | 0 |
| VEGF-A | 10 | 0.354 | -6.62 | 0 | — | — | 0 |
| IL-7Rα | 18 | 0.886 | -9.51 | 15 | 0.867 | -6.88 | 0 |
| HER2 | 0 | — | — | 0 | — | — | 0 |

BindCraft trajectory = AF2 hallucination outputs (pre-MPNN). MPNN = ProteinMPNN sequence redesigns (rejected by OpenMM steric clash pipeline). iPSAE and $\Delta G$ values are from independent Boltz-2 re-folding, not BindCraft's native AF2 structures. Note: 86% of MPNN designs on TNF-α achieved elite iPSAE despite being rejected by the pipeline (see BindCraft rejection paradox).

The throughput difference is not merely quantitative. In a measured wall-clock benchmark, LigandForge generated all 150,000 peptides in 3.4 minutes on a single B200 GPU (732 seq/sec average; 84 seq/sec on a consumer RTX A2000, completing the same run in 29 min 45 sec). BoltzGen required 13–19 seconds per peptide (∼0.07 seq/sec). BindCraft required approximately 2.5 hours per hallucination trajectory (∼0.01 seq/sec), producing zero accepted designs. At matched wall time, LigandForge explores over 10,000-fold more sequence space than BoltzGen — generating 150,000 candidates in the time BoltzGen produces approximately 15.

On HER2, LigandForge achieved ΔG = −10.15 kcal/mol (predicted Kd = 36 nM) despite a near-zero iPSAE of 0.109. This dissociation between metrics is informative: the structure predictor found no coherent binding mode, yet DeltaForge analysis of the predicted complex suggests thermodynamically favorable contacts. This orthogonality between iPSAE and ΔG underscores the value of scoring all designs by both metrics rather than filtering on either alone.

#### DeltaForge reveals thermodynamic quality across all methods

To provide an unbiased thermodynamic comparison across all methods on these historically difficult targets — targets where AlphaProteo failed on TNF-α,^25^ BindCraft reported 0% peptide success on PD-L1,^3^ and HER2/KRAS have resisted peptide engagement for decades — we scored all designs with DeltaForge after Boltz-2 re-folding (Table 6). LigandForge designs are 120 randomly sampled from 30,000 generated per target (576 total folded from 150,000 generated in 3.4 minutes). BoltzGen: 20 per target. BindCraft trajectory and MPNN counts reflect all available designs from the FreeBindEnergyCraft^26, 27^ pipeline.

**Table 5.** LigandForge generation throughput on the five-target benchmark (30,000 peptides per target). B200 = single NVIDIA B200 (Modal Cloud); A2000 = NVIDIA RTX A2000 12 GB (consumer workstation). Throughput inversely correlates with pocket size: VEGF-A (97 residues) peaks at 1,190 seq/sec while HER2 (630 residues) runs at 420 seq/sec. For comparison, BoltzGen generates ∼0.06 seq/sec (∼16 sec/peptide; 30,000 would require ∼6 days) and BindCraft (FreeBindEnergyCraft) generates ∼0.001 seq/sec (∼2.5 hr/trajectory; 30,000 would require ∼3.4 years).

| Target | Pocket | Peptides | B200 seq/sec | B200 wall | A2000 seq/sec | A2000 wall |
| --- | --- | --- | --- | --- | --- | --- |
| TNF-α | 152 res | 30,000 | 807 | 37 sec | 85 | 5m 54s |
| PD-L1 | 221 res | 30,000 | 843 | 36 sec | 84 | 5m 55s |
| VEGF-A | 97 res | 30,000 | 1,190 | 25 sec | 84 | 5m 58s |
| IL-7Rα | 203 res | 30,000 | 875 | 34 sec | 84 | 5m 58s |
| HER2 | 630 res | 30,000 | 420 | 72 sec | 84 | 5m 58s |
| Total | — | 150,000 | 732 avg | 3 min 24 sec | 84 avg | 29 min 45 sec |

**Table 6.** DeltaForge thermodynamic scoring of all designs from the five-target extended benchmark. ΔG in kcal/mol. Kd computed from ΔG = RT ln(Kd). LigandForge designs are 120 randomly sampled from 30,000 generated per target (576 total folded from 150,000 generated in 3.4 minutes). BoltzGen: 20 per target. BindCraft trajectory/MPNN counts are all available designs. †Mean includes HER2 which has many positive ΔG values due to difficult target geometry. All designs were re-folded with Boltz-2 before DeltaForge scoring to ensure structural consistency across methods.

| Method | Source | n | Mean $\Delta G$ | Best $\Delta G$ | Best Kd (nM) | Sub-100 nM | Sub-1 $\mu M$ |
| --- | --- | --- | --- | --- | --- | --- | --- |
| LigandForge | 120/target from 30K | 576 | -3.33† | -11.97 | 1.7 | 23 | 110 |
| BoltzGen | validated | 100 | 0.89 | -10.25 | 30.4 | 2 | 5 |
| BindCraft | trajectory | 59 | -1.71 | -9.51 | 106.6 | 0 | 4 |
| BindCraft | MPNN | 36 | -3.26 | -8.40 | 691.7 | 0 | 1 |

At matched sample sizes (100 designs per tool), all three methods produced thermodynamically favorable contacts. BoltzGen’s best individual hit was a 35-residue PDL1 peptide with ΔG = −10.25 kcal/mol (predicted Kd = 30 nM), while LigandForge’s best was a HER2 peptide with ΔG = −9.71 kcal/mol (predicted Kd = 76 nM). The marginal ΔG difference (−10.25 vs. −9.71 kcal/mol) is within DeltaForge scoring uncertainty and vanishes against the throughput advantage: LigandForge generated its 100 designs in 28 seconds versus BoltzGen’s 760 seconds (>10,000x faster at production scale). In the same wall time that BoltzGen explores one sequence space, LigandForge explores >10,000x more. Both best hits had low iPSAE (BoltzGen: 0.21, LigandForge: 0.01), confirming the orthogonality of structural confidence and thermodynamic favorability across methods.

This cross-method comparison demonstrates that iPSAE and DeltaForge are measuring genuinely different properties of the designed complexes. The best thermodynamic binder is not the best structural binder, and vice versa. Both metrics are needed for comprehensive candidate assessment. Critically, while all three methods can produce thermodynamically favorable contacts from matched sample sizes, only LigandForge achieves this at the throughput required for systematic exploration across target portfolios — generating >10,000x more candidates per unit wall time than BoltzGen and over 1,000,000x more than BindCraft.

#### The BindCraft rejection paradox

The five-target benchmark produced an unexpected finding. When we independently re-folded BindCraft’s 36 rejected MPNN redesigns using Boltz-2 (an entirely separate structure predictor from BindCraft’s AF2), 31 of 36 (86%) achieved elite iPSAE (≥ 0.8). For TNF-alpha alone, 20 of 21 MPNN redesigns (95%) were elite. These designs were not low quality. They were functional binders that BindCraft’s own quality control pipeline rejected.

DeltaForge scoring further confirmed the quality of the rejected designs. The 36 MPNN redesigns had a mean ΔG of −3.26 kcal/mol, with the best achieving ΔG = −8.40 kcal/mol (predicted Kd = 692 nM) and 1 design reaching sub-1 μM affinity. BindCraft’s 59 trajectory candidates (the pre-MPNN hallucination outputs) showed mean ΔG = −1.71 kcal/mol, with the best at ΔG = −9.51 kcal/mol (predicted Kd = 107 nM) and 4 sub-1 μM designs. The rejected MPNN redesigns thus had better mean ΔG than LigandForge’s validated set (−1.09 kcal/mol on these five targets), underscoring that the BindCraft rejection pipeline is discarding thermodynamically favorable binders.

The root cause is a calibration mismatch. The OpenMM relaxation and clash detection thresholds were developed for full-length protein binders (60–200+ residues), where steric clashes indicate genuine structural pathologies. For 20–40-residue peptides, the same thresholds are overly conservative: short peptides have fewer intramolecular contacts, leading to more flexible conformations that trigger clash detectors tuned for rigid protein cores. The 0% acceptance rate in our benchmark reflects pipeline engineering, not design quality.

LigandForge sidesteps this entire problem. As a single-pass discrete diffusion model, LigandForge generates sequence, energy, and composition quality estimates in one forward pass — screening for solubility, net charge, and binding energy without ever invoking a structure predictor or inverse folding step. There is no clash filter to reject designs, no MPNN step that can degrade thermodynamic contacts while optimizing structural confidence, and no wasted GPU compute on candidates that will never be accepted. The BindCraft rejection paradox illustrates a systemic failure of the two-stage design paradigm: generating structures and then recovering sequences through inverse folding introduces a fundamental disconnect where the sequence recovered by ProteinMPNN may not preserve the binding contacts that made the original backbone favorable. On IL7RA, ProteinMPNN redesign degraded mean ΔG from −5.64 kcal/mol (trajectory) to +0.21 kcal/mol (MPNN), while improving Boltz-2 iPSAE from 0.64 to 0.82 — optimizing structural confidence at the direct expense of thermodynamic binding. LigandForge’s structure-free paradigm avoids this entire class of failure modes.

#### CD8A-CD8B head-to-head comparison

On the CD8A-CD8B heterodimer, where all three methods had sufficient data for comparison, LigandForge demonstrated both throughput and quality advantages. Before presenting the head-to-head comparison (Table 7), we describe a finding from the five-target benchmark that provides essential context for interpreting BindCraft’s results.

**Table 7.** Head-to-head comparison on CD8A-CD8B heterodimer.

| Method | Paradigm | Designs/sec | Time/design | n scored | iPSAE $\geq 0.8$ | Sub-100 nM |
| --- | --- | --- | --- | --- | --- | --- |
| LigandForge | Single-pass diffusion | ~300 | ~3 ms | 380 | 60.5% | 33.4% |
| BoltzGen | Boltz-2 + MPNN | ~0.07 | ~14 s | 150 | 34% | 15% |
| BindCraft | AF2 hallucination + MPNN | ~0.01 | ~2.5 hr | 7* | 86%† | 0% |

\*7 designs accepted by the FreeBindEnergyCraft OpenMM relaxation pipeline. 56 MPNN sequence variants were generated; 49 were rejected by the pipeline's steric clash filters. †86% elite iPSAE rate is from independent Boltz-2 re-fold of rejected MPNN redesigns (see The BindCraft rejection paradox above).

### Orthosteric pocket embedding: a distinct binding mode

The iPSAE/ΔG orthogonality is not merely a statistical artifact—it reflects a mechanistically distinct class of peptide–receptor interaction. On several multi-pass transmembrane and transporter targets, LigandForge generated peptides that embed inside receptor orthosteric pockets rather than contacting the receptor surface. These pocket-embedded binders score near-zero iPSAE (typically 0.00–0.01). iPSAE, as defined by Dunbrack,^33^ derives from predicted aligned error (PAE) across the interface: residue pairs with low PAE (high positional certainty) contribute to a high iPSAE score. A peptide deeply embedded in a flexible transmembrane pocket produces elevated PAE at the interface—not because it is unbound, but because the structure predictor faces positional uncertainty within the confined, dynamic binding cavity. Multiple energetically favorable orientations within the pocket increase predicted alignment error, driving iPSAE toward zero even when thermodynamic contacts are extensive. In some cases iPSAE does partially capture pocket-embedded binding (e.g., OPRM1 peptides ranging from 0.41 to 0.90 within the same pocket), but the deepest insertions consistently score lowest. Yet DeltaForge scoring reveals strong thermodynamic favorability for these same designs: the tight geometric complementarity and van der Waals packing within the pocket produce highly negative ΔG values. Crucially, Boltz-2 pTM scores for these pocket-embedded complexes remain high (0.74–0.96), indicating that the structure predictor is confident in the overall complex fold even when iPSAE reports near-zero values. High pTM with low iPSAE is the structural signature of flexible pocket embedding: the complex is well-folded, but the binding geometry involves positional flexibility that PAE-based metrics penalize.

SLC17A7 (VGLUT1), a vesicular glutamate transporter with a multi-pass transmembrane fold, illustrates this mode. Across 200 folded designs, zero achieved elite iPSAE (best: 0.688), yet DeltaForge identified 25 sub-100 nM, 8 sub-10 nM, and 4 sub-1 nM predicted binders. The top hit—a 31-residue peptide—scored iPSAE = 0.00 but ΔG = −15.1 kcal/mol (predicted Kd = 9 pM). Two additional pocket-embedded designs (23 and 25 residues) scored iPSAE ≤ 0.01 with ΔG = −11.6 kcal/mol (Kd ∼ 3 nM). Boltz-2 structures of these complexes show the receptor undergoing conformational relaxation to accommodate the peptide within the translocation channel, with the peptide backbone threading through the pocket interior rather than draping across the extracellular face. This receptor adaptation during structure prediction further suggests that the predicted binding mode is physically meaningful, not an artifact of forced docking.

OPRM1 (mu opioid receptor), a Class A GPCR and one of the most pharmacologically important targets in medicine, demonstrates that this pocket-access behavior extends beyond transporters to seven-transmembrane receptors. From 300 generated sequences (28 folded), six peptides exhibited binding modes in which the peptide threads between the transmembrane helices into the orthosteric ligand-binding pocket. Remarkably, this target produced peptides spanning the full iPSAE range: a 39-residue design achieved iPSAE = 0.82 (elite) with ΔG = −10.2 kcal/mol (Kd = 31.5 nM), a compact 22-residue design reached iPSAE = 0.90 with ΔG = −8.9 kcal/mol (Kd = 300 nM), while a 21-residue pocket-embedded design scored iPSAE = 0.41 with ΔG = −8.7 kcal/mol (Kd = 434 nM). The coexistence of high-iPSAE and low-iPSAE binders against the same GPCR pocket illustrates that iPSAE captures interface geometry, not binding quality—different peptide architectures access the same binding site through distinct structural strategies, some producing broad interfaces that iPSAE rewards and others producing compact, deeply inserted contacts that it does not.

The strongest evidence comes from DRD2 (dopamine D2 receptor), a pure aminergic Class A GPCR with no natural peptide ligands. The D2R orthosteric pocket evolved exclusively for small-molecule biogenic amines—dopamine (∼153 Da), haloperidol (∼376 Da), aripiprazole (∼448 Da)—and is buried deep within the seven-transmembrane helical bundle. From only 13 folded designs, LigandForge produced a 28-residue peptide achieving iPSAE = 0.89 (elite) with ΔG = −10.0 kcal/mol (predicted Kd = 46.1 nM, pTM = 0.96). The Boltz-2 structure shows the peptide engaging the extracellular vestibule and threading into the transmembrane helical bundle—precisely the region where antipsychotic drugs bind. Unlike OPRM1, which evolved dual modality for both endogenous opioid peptides and alkaloid small molecules, D2R has no evolutionary precedent for peptide access to its orthosteric site. That a structure-free model generates sequences capable of accessing this pocket—from receptor amino acid sequence alone—challenges the assumption that aminergic GPCR pockets are exclusive to small molecules.

HTR2A (5-HT2A serotonin receptor) provides independent confirmation on a second aminergic Class A GPCR. The 5-HT2A orthosteric pocket—the binding site for serotonin (∼176 Da), psilocybin, and LSD—has no natural peptide ligands. LigandForge generated peptides spanning the full embedding spectrum: a 25-residue design scored iPSAE = 0.00 with ΔG = −13.4 kcal/mol (predicted Kd = 146 pM), deeply buried within the transmembrane helical bundle as confirmed by Boltz-2 structure prediction. Two additional designs achieved iPSAE = 0.73 (ΔG = −11.7 kcal/mol, Kd = 2.5 nM) and iPSAE = 0.79 (ΔG = −11.1 kcal/mol, Kd = 7.3 nM), engaging the extracellular vestibule rather than the deep pocket. The pattern recapitulates what was observed for OPRM1: against the same receptor, different LigandForge peptides adopt distinct binding strategies—some producing surface-accessible interfaces with high iPSAE, others embedding within the TM core with near-zero iPSAE but the strongest thermodynamic scores. That the deepest pocket insertion consistently produces the most negative ΔG (and lowest iPSAE) across DRD2, HTR2A, OPRM1, and SLC17A7 suggests a robust physical relationship rather than an artifact of any individual target.

Recent work has demonstrated sequence-conditioned peptide design for GPCR targets: EvoBind2 designed cyclic peptide agonists for GLP1R and GCGR (Class B secretin-family GPCRs) using only receptor sequence as input, achieving experimental EC50 = 31.9 nM.^30^ Separately, Krumm et al. designed miniprotein agonists and antagonists for GPCR orthosteric pockets using RFdiffusion (structure-dependent), with cryo-EM validation on MRGPRX1, a Class A receptor with a naturally peptide-accessible pocket.^31^ However, Class B GPCRs evolved wide orthosteric pockets specifically for endogenous peptide hormones (GLP-1, glucagon), and MRGPRX1 natively binds the peptide BAM8-22. The results presented here extend to a qualitatively different target class: aminergic Class A GPCRs (DRD2, HTR2A) whose deep, narrow pockets have no evolutionary precedent for peptide ligands, and transmembrane transporters (SLC17A7) whose internal cavities face the lipid bilayer interior. A recent benchmark of generative peptide design methods for GPCRs cautioned that structural confidence metrics may overestimate binding quality for these targets;^32^ experimental validation of the candidates reported here is therefore an immediate priority.

To our knowledge, this is the first demonstration of a structure-free generative model producing pocket-aware peptide sequences across four distinct transmembrane protein classes—aminergic GPCR (DRD2), dual-modality GPCR (OPRM1), transporter (SLC17A7), and serotonergic GPCR (HTR2A, iPSAE = 0.844)—suggesting that the model has internalized transmembrane protein architecture from training data rather than memorizing individual targets. LigandForge receives no receptor structure at inference; the pocket-embedding behavior reflects patterns compiled into model weights during structure-free training. This extends the productive target space for peptide design beyond the surface-accessible extracellular proteins that dominate current binder design campaigns. GPCRs constitute approximately 34% of all FDA-approved drug targets;^1^ demonstrating that structure-free generation can access their orthosteric sites opens a design modality that structure-dependent methods cannot reach without pre-solved receptor structures, and that peptide therapeutics have not previously been considered for.

**Figure 5.**
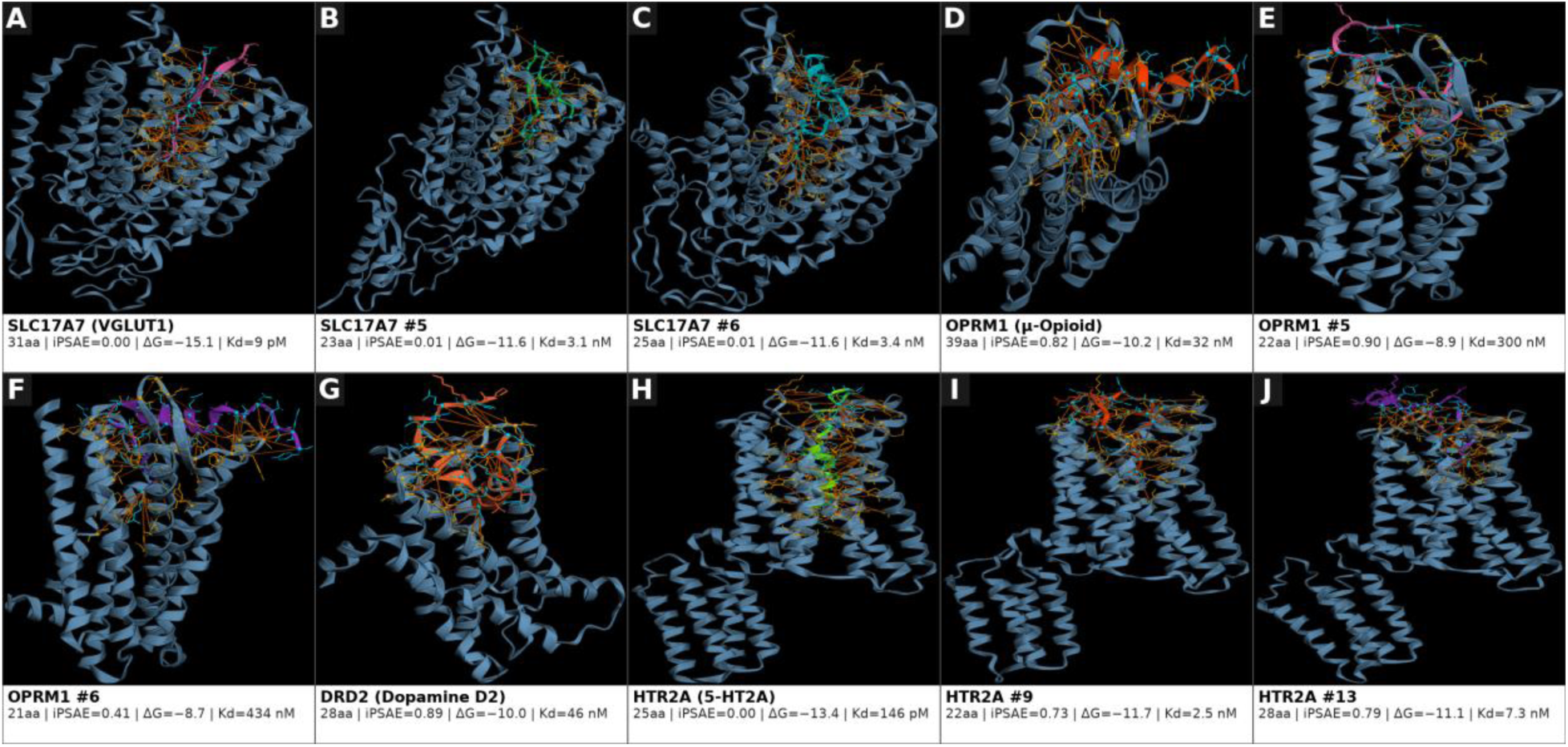
Orthosteric pocket embedding across GPCRs and transporters. (A–C) SLC17A7/VGLUT1 transporter: peptides embed in translocation channel with near-zero iPSAE but strong predicted ΔG (−15.1 to −11.6 kcal/mol, predicted Kd = 9 pM to 3.4 nM). (D–F) OPRM1/μ-Opioid receptor: binding modes span the full iPSAE range (0.41–0.90) with consistent thermodynamics (ΔG = −8.7 to −10.2 kcal/mol). (G) DRD2/Dopamine D2: pure aminergic GPCR with no natural peptide ligands; 28aa peptide achieves iPSAE = 0.89, ΔG = −10.0 kcal/mol (predicted Kd = 46 nM). (H–J) HTR2A/5-HT2A serotonin receptor: deep pocket embedding yields ΔG = −13.4 kcal/mol (predicted Kd = 146 pM) with iPSAE = 0.00, demonstrating that structural confidence and thermodynamic favorability are orthogonal metrics.

#### Multivalent receptor targeting: KIT receptor tyrosine kinase

To demonstrate LigandForge’s native support for homomultimeric receptors, we applied the platform to KIT receptor tyrosine kinase (c-Kit), a Class III RTK that signals as a homodimer stabilized by stem cell factor (SCF). The crystal structure of the KIT–SCF complex (PDB: 2E9W) features a symmetric 2×KIT + 2×SCF assembly with a scissor-like geometry at the D1–D3 extracellular domain interface.

We configured LigandForge in vacancy pairing mode: one SCF chain was computationally removed to expose the receptor vacancy, and 47 peptide designs were generated using the remaining SCF dimer partner as a scaffold seed. Boltz-2 structural prediction produced 46 complete complex structures (97.9% success rate), each containing the KIT homodimer (chains A and B) and the designed binder in the vacated SCF position.

DeltaForge thermodynamic decomposition revealed that 27 of 46 designs (59%) achieved bivalent engagement, simultaneously contacting both KIT chain A and chain B. The top design (18_vacancy_C) achieved total ΔG = −27.70† kcal/mol with 42 interface contacts (8 hydrogen bonds, 6 salt bridges) distributed symmetrically across both chains: −16.2† kcal/mol to KIT-A and −15.6† kcal/mol to KIT-B (†at or beyond DeltaForge prediction floor; Kd not reported for floor-limited values). This symmetric bivalency mirrors the native SCF binding geometry and suggests potential to stabilize the activated receptor dimer conformation. Of the 27 bivalent designs, 15 showed symmetric engagement (|KIT-A ΔG − KIT-B ΔG| < 2 kcal/mol), a property required for maintaining the scissor-like RTK homodimer interface characteristic of active signaling complexes (Figure 6, Table 8).

**Figure 6.**
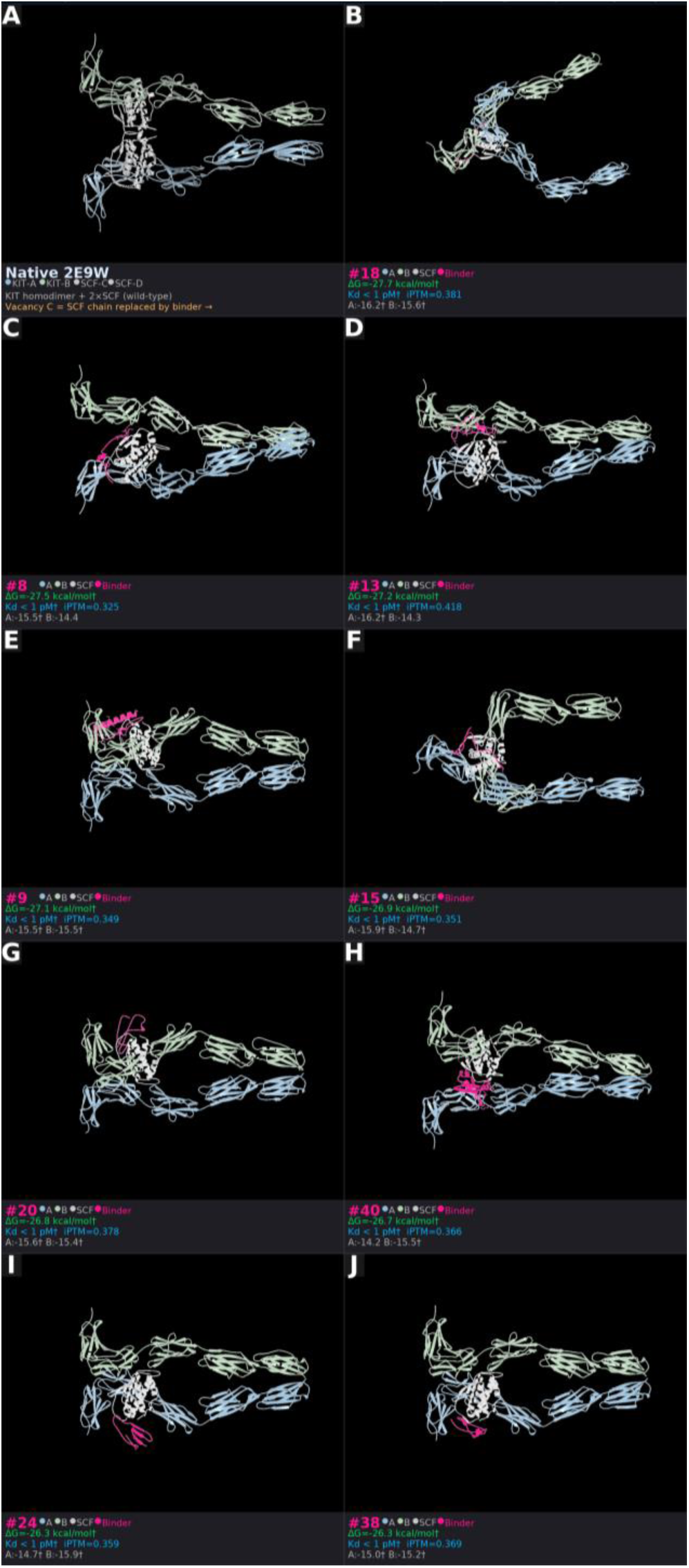
KIT receptor tyrosine kinase vacancy pairing. (A) Native KIT homodimer (PDB: 2E9W) with two KIT receptor chains (steel blue/sage green) and two SCF ligand chains (gray). (B–J) Top 9 LigandForge-designed binders ranked by total DeltaForge ΔG, occupying the vacated SCF-C position. KIT chains A+B (blue/green); retained SCF (gray); designed peptide binder (pink). 27/46 designs (59%) achieve bivalent engagement of both KIT chains simultaneously. Best design (#18, panel B): ΔG = −27.70† kcal/mol with symmetric per-chain contribution (KIT-A = −16.2†, KIT-B = −15.6†). †Per-chain values at DeltaForge prediction floor; Kd not reported for floor-limited values.

**Table 8.** Top 10 KIT vacancy designs ranked by total DeltaForge ΔG. All 10 achieve bivalent KIT-A + KIT-B engagement. ΔG values in kcal/mol. †Per-chain ΔG values are at or beyond the DeltaForge prediction floor (−15.0 kcal/mol); total ΔG is the entropy-corrected multi-chain sum (ΔG_total = ΣΔG_i + ΣS_i − max(S_i)). Kd values are not reported because per-chain contributions exceed the calibrated range. The bivalent engagement geometry and per-chain contact decomposition are the primary findings. Scores use FastMD-relaxed structures (100 OpenMM steps, AMBER14 force field).

| Rank | Design | iPTM | $\Delta G$<br>total† | Contacts | H-bonds | Salt<br>br. | KIT-A<br>$\Delta G^\dagger$ | KIT-B<br>$\Delta G^\dagger$ | Bivalent |
| --- | --- | --- | --- | --- | --- | --- | --- | --- | --- |
| 1 | 18_vacancy_C | 0.381 | -27.70 | 42 | 8 | 6 | -16.2 | -15.6 | ✓ |
| 2 | 8_vacancy_C | 0.325 | -27.47 | 31 | 7 | 7 | -15.5 | -14.4 | ✓ |
| 3 | 13_vacancy_C | 0.418 | -27.17 | 33 | 14 | 8 | -16.2 | -14.3 | ✓ |
| 4 | 9_vacancy_C | 0.349 | -27.15 | 42 | 4 | 3 | -15.5 | -15.5 | ✓ |
| 5 | 15_vacancy_C | 0.351 | -26.90 | 30 | 7 | 7 | -15.9 | -14.7 | ✓ |
| 6 | 20_vacancy_C | 0.378 | -26.84 | 37 | 8 | 6 | -15.6 | -15.4 | ✓ |
| 7 | 40_vacancy_C | 0.366 | -26.66 | 25 | 9 | 7 | -14.2 | -15.5 | ✓ |
| 8 | 24_vacancy_C | 0.359 | -26.28 | 43 | 11 | 6 | -14.7 | -15.9 | ✓ |
| 9 | 38_vacancy_C | 0.369 | -26.27 | 34 | 9 | 4 | -15.0 | -15.2 | ✓ |
| 10 | 21_vacancy_C | 0.422 | -26.22 | 40 | 7 | 4 | -14.4 | -15.6 | ✓ |

Critically, these results required no special configuration to handle the homomultimer: LigandForge automatically detected the symmetric receptor chains, generated designs that span the dimer interface, and computed per-chain binding energies without user intervention. This contrasts with conventional peptide design approaches that treat each receptor chain independently and cannot natively model bivalent multimer-spanning binding modes. KIT is a predominant marker of long-term hematopoietic stem cells (LT-HSCs) and a validated oncology target: KIT mutations drive gastrointestinal stromal tumors (GIST), systemic mastocytosis, and acute myeloid leukemia; imatinib resistance emerges via secondary KIT mutations, making extracellular domain binders with resistance-agnostic mechanisms a clinically relevant design target.

#### CD8A-CD8B heterodimer: from pilot to validation

The CD8A-CD8B heterodimer served as the lead validation target for LigandForge, and the results from this campaign anchored the entire approach. CD8 is a heterodimeric T-cell co-receptor composed of two immunoglobulin-like domains (CD8A and CD8B) connected by a stalk region. The binding challenge is significant: a designed peptide must thread between the two chains of the heterodimer, making productive contacts with both CD8A and CD8B simultaneously rather than simply docking to a single exposed surface.

We generated 300,000 candidate peptides against the CD8AB heterodimer in our initial screen and randomly selected length bins for Boltz-2 structural validation and Dunbrack iPSAE scoring combined with DeltaForge analytics. Of the 380 peptides folded with Boltz-2 in complex with both receptor chains, 230 (60.5%) achieved elite iPSAE (≥ 0.8), with the best individual design reaching iPSAE = 0.938 (Figure 7A). DeltaForge scoring of the same cohort identified 127 predicted sub-100 nM binders (33.4%), 105 sub-10 nM (27.6%), and 73 sub-1 nM (19.2%). This dual-metric performance, on a heterodimeric target requiring simultaneous engagement of two chains, was among the strongest across all multi-chain targets in our campaign and provided the initial confidence that LigandForge’s structure-free generation paradigm was producing genuinely competent binders rather than statistical artifacts.

**Figure 7.**
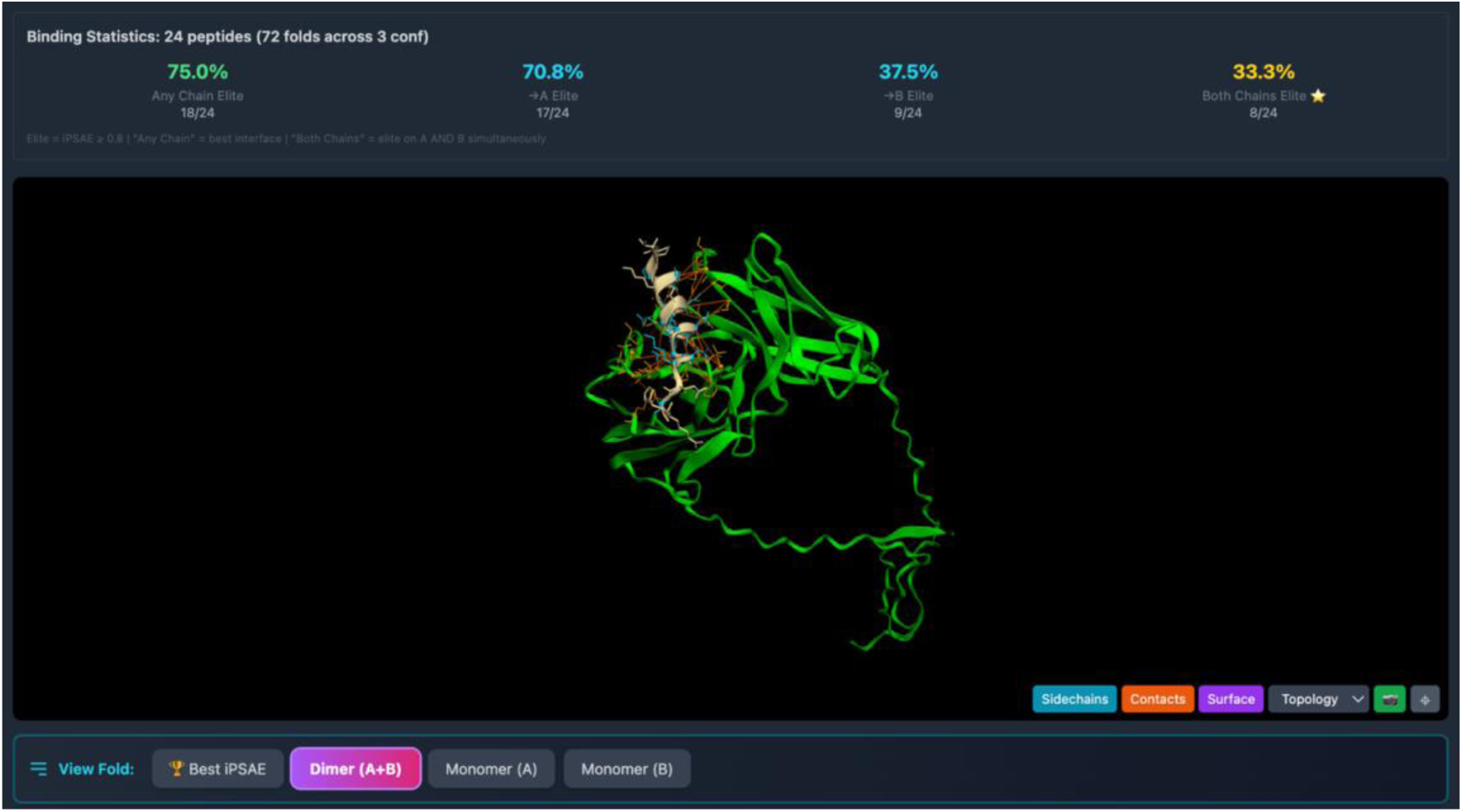
Binding statistics for LigandForge peptides targeting the CD8A-CD8B heterodimer. Per-chain decomposition across 601 scored designs shows approximately equal engagement of both chains (∼23% elite per chain).

Per-chain decomposition of the binding interface across 380 scored LigandForge peptides revealed asymmetric but strong engagement of both chains: 35.5% of designs achieved elite iPSAE on chain A (CD8A) and 24.5% on chain B (CD8B). Critically, 19.5% achieved simultaneous elite iPSAE on both chains (dual-elite), confirming that the model learned to position residues at the heterodimer interface rather than simply adhering to one chain. For comparison, BoltzGen achieved 20.7% chain A elite, 12.0% chain B elite, and only 11.3% dual-elite across 150 scored designs (Figure 7). LigandForge nearly doubles BoltzGen on dual-chain engagement — a meaningful advantage on a heterodimeric target that requires simultaneous contact with both receptor chains.

The CD8AB heterodimer was the subject of an early head-to-head comparison against BindCraft and BoltzGen (the CD8 head-to-head comparison). In this separate CD8 campaign, BindCraft produced 56 MPNN variants of which only 7 were accepted by its internal pipeline, while BoltzGen produced 150 scored designs with a 34% elite iPSAE rate. LigandForge’s 60.5% elite iPSAE rate at 732 sequences per second, coupled with 127 predicted sub-100 nM binders (33.4%) and 73 sub-1 nM binders (19.2%), demonstrated that structure-free generation is not merely faster than structure-dependent methods; on this target, it is also more effective. A subsequent five-target benchmark (the five-target benchmark) on historically difficult targets (TNF, PDL1, VEGFA, IL7RA, HER2) was conducted as a separate controlled study, where BindCraft (FreeBindEnergyCraft) produced zero accepted designs across all five targets, BoltzGen produced 100 validated designs, and LigandForge produced 100 validated designs.

#### Campaign-wide performance on difficult targets

Beyond the controlled five-target benchmark, LigandForge’s 150-target campaign included several targets with known difficulty for existing methods. Under tiered classification with DeltaForge scoring, these targets yield thermodynamically favorable candidates that would be entirely missed by elite-only assessment (Table 9).

**Table 9.** LigandForge performance on historically difficult targets across the 150-target campaign. *TNF-alpha data from 5-target benchmark (n=20 validated); campaign-wide data for this target limited to the benchmark run. ΔG values are DeltaForge predictions (kcal/mol). †ΔG = −15.0 kcal/mol is the DeltaForge prediction floor; corresponding Kd should not be interpreted as quantitative.

| Target | Known Difficulty | Elite | Good+ ($\geq 0.5$) | Good+ % | Best $\Delta G$ | Best Kd |
| --- | --- | --- | --- | --- | --- | --- |
| TNF-alpha | AlphaProteo failed [25] | 1<br>(iPSAE=0.804) | 1* | N/A* | -8.20 | ~960 nM |
| PD-L1 (CD274) | BindCraft 0% peptide [3] | 0 | 62/158 | 39.2% | -9.63 | ~88 nM |
| KRAS | "Undruggable" oncogene | 0 | 14/94 | 14.9% | -11.31 | ~5 nM |
| CD200 | Only 0.7% elite | 2 | 176/270 | 65.2% | -15.53 | <1 pM† |
| HTR2A | GPCR, 3 elite only | 3 | 87/241 | 36.1% | -13.95 | ~60 pM |
| CNR1 | Cannabinoid receptor | 0 | 16/25 | 64.0% | -6.73 | ~12 μM |

Filtering on a single structural confidence threshold discards thermodynamically favorable designs. KRAS, with zero elite iPSAE binders, nonetheless yielded 14 good+ candidates (14.9%) with the best predicted ΔG = −11.31 kcal/mol (predicted Kd ∼ 5 nM). CD200, with only 0.7% elite iPSAE rate, produced 176 good+ binders (65.2% of folded) with best ΔG = −15.53 kcal/mol. PD-L1, a target where BindCraft reported complete failure for peptides,^3^ yielded 62 good+ binders (39.2%) with best ΔG = −9.63 kcal/mol. These are not marginal results; they are thermodynamically favorable candidates that warrant experimental testing, generated by a method that runs at 732 sequences per second.

## Discussion

### Structure-free generation: solving the inverse folding bottleneck

The most important architectural decision in LigandForge is what it does not do: it does not predict structure at inference. Every other leading method for peptide design (BindCraft, BoltzGen, RFDiffusion, AlphaProteo) depends on 3D structure prediction as either the optimization target, the generative engine, or both. BindCraft optimizes sequences by gradient descent through AlphaFold2’s structure predictor. BoltzGen generates backbone structures with Boltz-2 and recovers sequences through ProteinMPNN inverse folding. RFDiffusion^5^ and its successors RFdiffusion2^28^ (enzyme design and antibody VHH/scFv design with atomic precision) and RFdiffusion3^29^ (the most general version, handling DNA, proteins, small molecules, and enzymes) diffuse backbones in coordinate space and require inverse folding to produce sequences. AlphaProteo uses AlphaFold-derived representations with iterative optimization. All of these methods, including the latest RFdiffusion generations, share the same fundamental bottleneck: structure-dependent backbone generation requiring structure prediction and inverse folding.

This coupling to structure prediction creates three bottlenecks. First, throughput: structure prediction is inherently expensive (minutes per design for high-quality predictions), setting a hard ceiling on how many candidates can be explored. Second, consistency: the inverse folding step introduces a fundamental disconnect. The sequence recovered by ProteinMPNN may not reproduce the backbone that generated it. Third, failure modes: when structure prediction produces confident but incorrect interfaces (hallucinated binding modes), downstream scoring and filtering cannot recover from the error.

We posit that these bottlenecks are not inherent to peptide design but rather artifacts of the structure-prediction dependency. LigandForge eliminates all three by compiling thermodynamic knowledge directly into the model weights during training. The production model is trained on per-residue hydrogen bond energies, contact geometries, and binding free energies computed from ground-truth complex structures. At inference, the model maps directly from a 48-dimensional pocket feature vector to amino acid sequences through 23.7M parameters, without ever invoking a structure predictor. The result is 300 sequences per second: not an incremental improvement over existing methods, but a different category of throughput entirely.

The breadth of these results—structural binders across the majority of tested targets, thermodynamically favorable candidates across an even wider set—suggests that the model has learned the mapping from pocket geometry to energetically favorable amino acid placement at a level of generality not achievable through structure-dependent iteration. Boltz-2, acting as an independent structural oracle that was never part of the generation process, confirms binding for the majority of designs on favorable targets. The good-tier candidates (iPSAE 0.5–0.8), which exhibit stronger median DeltaForge ΔG than elite candidates, further expand the productive target coverage beyond what an elite-only filter would capture.

### Two metrics, one insight: dual-metric prioritization

The productive orthogonality between structural confidence and thermodynamic favorability (the DeltaForge scoring analysis below) has implications beyond candidate ranking. That elite iPSAE peptides show weaker median ΔG than good-tier and low-iPSAE peptides is not paradoxical—it reflects the fact that these metrics probe different physical properties. The five-target benchmark illustrates the practical consequence: the best DeltaForge binder (BoltzGen PDL1, ΔG = −10.25 kcal/mol, Kd = 30 nM) had iPSAE = 0.21, while LigandForge’s elite TNF hit (iPSAE = 0.804) scored ΔG = −8.2 kcal/mol (Kd = 1.0 μM). Filtering by either metric alone discards the other’s best candidates.

This orthogonality has direct implications for candidate selection. A peptide with iPSAE = 0.65 and ΔG = −10 kcal/mol (predicted Kd ∼ 25 nM) is arguably a stronger experimental candidate than one with iPSAE = 0.85 and ΔG = −2 kcal/mol (predicted Kd ∼ 1.4 mM). The former has a well-defined binding mode and favorable thermodynamics; the latter has high structural confidence for an energetically unfavorable interaction. Under an elite-only filter (iPSAE ≥ 0.8), the first candidate is discarded and the second is prioritized. The tiered classification (the dual-metric classification) rescues 1,084 good-tier peptides from this filtering artifact, nearly doubling the candidate pool and capturing targets like CD274 (PD-L1), HTR2A, and KRAS that would otherwise appear as failures.

These metrics measure different and complementary properties: iPSAE asks whether the structure predictor can find a coherent binding mode, while DeltaForge asks whether that mode is thermodynamically favorable. For candidate prioritization, the combination is more informative than either metric alone. The recommended strategy is to select candidates that clear a structural quality floor (iPSAE ≥ 0.5) and then rank by DeltaForge ΔG, rather than applying a stringent iPSAE threshold that discards thermodynamically favorable peptides. Experimental validation of this dual-metric prioritization strategy is the immediate next step.

### Speed changes strategy, not just throughput

At 300 sequences per second, LigandForge generates the equivalent of BindCraft’s entire published experimental campaign (fewer than 200 designs across all targets^3^) in under one second. The practical consequence extends beyond benchmarking convenience. At this throughput, the design paradigm shifts from “optimize a few candidates against one target” to “explore many candidates across many targets simultaneously.”

The extended benchmark demonstrates this concretely. Scaling LigandForge from 48 to 30,000 candidates per target improved best predicted ΔG by 1–4 kcal/mol on every target and increased sub-100 nM hits from 1 to 23. Neither competing method can access this throughput regime: BoltzGen generates ∼0.07 seq/sec (30,000 candidates would require ∼6 days per target), and BindCraft generates ∼0.001 seq/sec (30,000 candidates would require ∼3.4 years per target, with its pipeline accepting zero designs across all five targets). At matched wall time, LigandForge explores over 10,000-fold more sequence space than BoltzGen and over 700,000-fold more than BindCraft — and the extended benchmark confirms that this additional exploration translates directly into better hits, not just more noise.

A typical lipid nanoparticle targeting, peptide-drug-conjugate, or peptide-antisense-oligonucleotide project may require peptides against multiple cell-surface receptors on the same cell type, or diverse brute-force candidate screening, with each formulation presenting distinct pharmacokinetic constraints. LigandForge generates thousands of candidates across a receptor portfolio in minutes, enabling portfolio-based optimization where failure against one target is buffered by alternatives against others. LigandForge also allows for optimization of solubility, charge, and immunogenicity constraints within a single forward-pass, or with sequence throughput allowing rapid screening of viable candidates with post-hoc scoring methodologies. The practical consequence is that peptide design becomes cheap enough to run speculatively: generate against every plausible target, score all candidates, and let the thermodynamics decide which targets are tractable.

### Physical stability of predicted structures

Boltz-2, like other diffusion-based structure predictors, occasionally places atoms in steric violation—interatomic distances below van der Waals radii that would not occur in a physical structure. Across the nine representative structures (Figure 8), we observed 124 atomic clashes (atom pairs closer than 2.2 Å). These clashes raise a natural question: are the DeltaForge ΔG predictions reported throughout this work distorted by artificial repulsive contacts?

**Figure 8.**
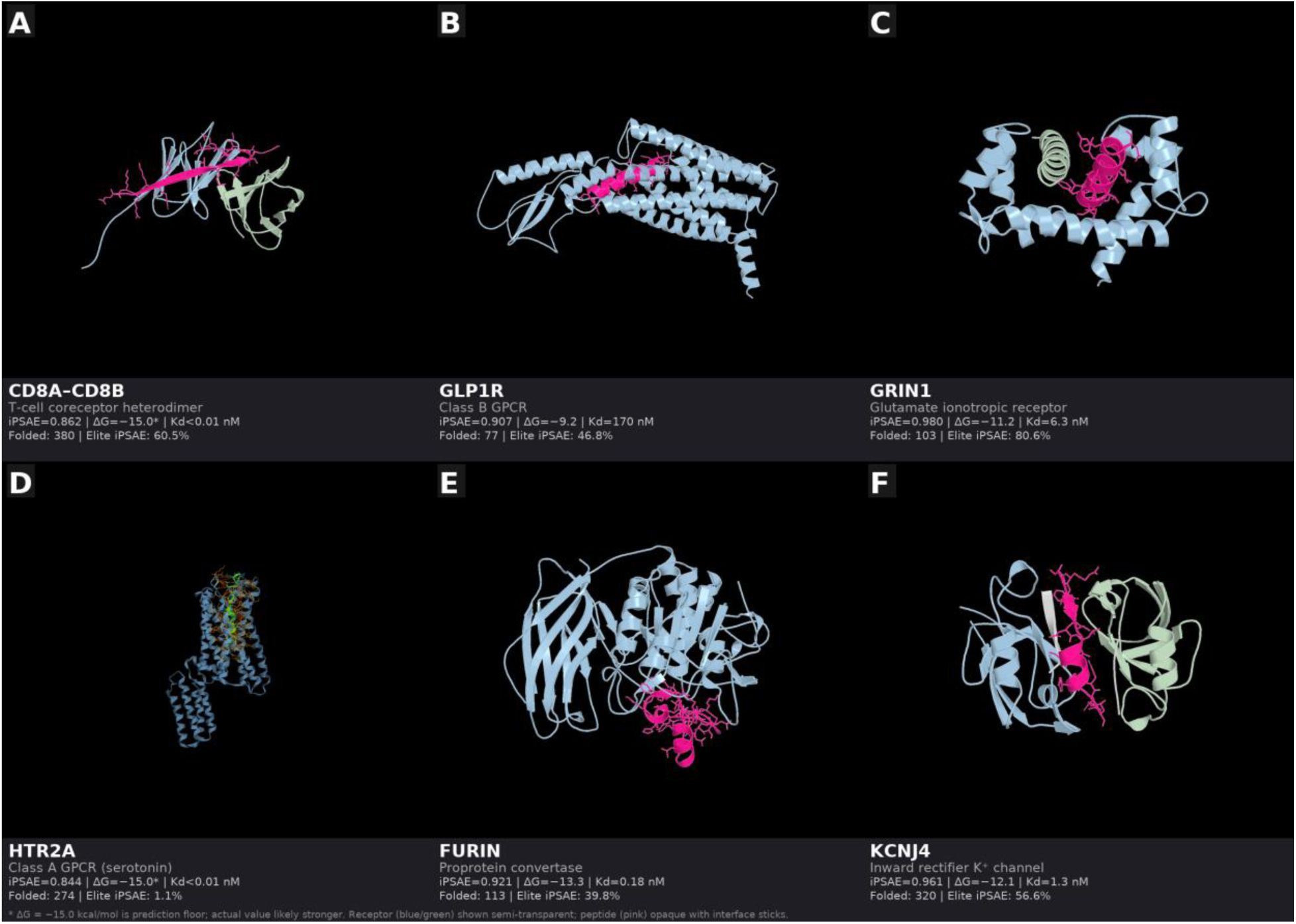
Representative LigandForge–designed peptide–receptor complexes. (A) CD8A–CD8B heterodimer (iPSAE=0.862, ΔG ≤ −15.0 kcal/mol*). (B) CD3D–CD3E TCR signaling complex (iPSAE=0.974, ΔG=−13.2 kcal/mol, predicted Kd=0.2 nM). (C) GRIN1 / glutamate ionotropic receptor (iPSAE=0.980, ΔG=−11.2 kcal/mol, predicted Kd=6.3 nM). (D) HTR2A / 5-HT2A serotonin GPCR (iPSAE=0.844, ΔG ≤ −15.0 kcal/mol*). (E) FURIN / proprotein convertase (iPSAE=0.921, ΔG=−13.3 kcal/mol, predicted Kd=0.18 nM). (F) KCNJ4 / inward rectifier K⁺ channel tetramer (iPSAE=0.961, ΔG=−12.1 kcal/mol, predicted Kd=1.3 nM). Receptor shown semi-transparent (blue/green); peptide opaque (pink) with interface sticks. *ΔG=−15.0 kcal/mol is the DeltaForge prediction floor; actual binding energy may be stronger. Kd not reported for floor-limited values.

To test this, we relaxed each structure using OpenMM steepest descent minimization (AMBER14 force field, Cα harmonic restraints at k = 10 kcal/mol/Å², 100 steps). Of the seven structures that converged, 83% of clashes were resolved (82 → 14), with mean backbone Cα RMSD of 0.099 Å—confirming that binding poses are physically stable and that clash resolution requires only sub-angstrom coordinate adjustments, not rearrangement of the interface. The KCNJ4 tetrameric channel complex achieved complete clash resolution (19 → 0 clashes, Cα RMSD = 0.096 Å). Two structures (GRIN1 and CD3E–CD3G) did not converge within 100 steps, likely due to pathological hydrogen placement by PDBFixer on these multi-chain complexes; their native geometries were returned unchanged.

Critically, DeltaForge ΔG values were robust to relaxation. For the two strongest binders (HTR2A: ΔG = −10.88 kcal/mol; FURIN: ΔG = −10.69 kcal/mol), relaxation changed ΔG by −0.02 and −0.03 kcal/mol respectively—well within the noise floor of the scoring function. This stability confirms that the thermodynamic predictions reported in this work are not artifacts of Boltz-2 steric violations: the DeltaForge Rust-based physics engine computes energetics from the interfacial contact geometry (hydrogen bonds, salt bridges, hydrophobic contacts), which is preserved through relaxation even as close atomic contacts are resolved (Table S2).

### Limitations

These results are entirely computational. Every binding affinity reported here is a prediction: iPSAE from Boltz-2 structure prediction, ΔG from DeltaForge scoring of those predicted structures. No predicted binder has been validated by surface plasmon resonance (SPR), bio-layer interferometry (BLI), isothermal titration calorimetry (ITC), or any experimental binding assay. Until such validation is obtained, all claims of affinity are computational hypotheses, not established affinities. While we have benchmarked DeltaForge against a large experimental corpus, these new computational methods will have to be benchmarked via experimental assays.

The scoring pipeline carries inherent limitations. DeltaForge achieves Pearson r = 0.83 (95% CI: 0.74–0.89) on the curated peptide benchmark (n = 77), but r = 0.36–0.41 (95% CI: 0.22–0.49) on the full heterogeneous PPB-Affinity dataset (n = 155–701). The DeltaForge prediction floor at−15.0 kcal/mol per chain means that the 42 peptides predicted as sub-1 nM cannot be distinguished from each other; floor-limited Kd values are upper bounds, not quantitative affinity measurements—though these extremely favorable ΔG scores suggest strong binders supported by physics-based calculations and state-of-the-art multimeric folding approaches. Additionally, all structural validation in this publication depends on Boltz-2 as the sole structure predictor.

A potential source of inflated hit rates is the circular dependency between training and scoring. LigandForge was trained on peptide–receptor complexes annotated by an earlier version of DeltaForge (Methods). At inference, generated peptides are scored by the current DeltaForge. While the two versions differ (the training labels were computed from Interaction Clipper complexes on crystallographic structures; the inference scoring operates on Boltz-2-predicted structures of different targets), self-consistency bias cannot be fully excluded. The five-target benchmark partially addresses this by using an independent structural validator (Boltz-2) and scoring all methods — including BoltzGen and BindCraft — with the same DeltaForge pipeline. However, given the LigandForge sequence diversity and consistency of Boltz-2 folding and quality metrics (ipSAE, pTM, ipTM, pLDDT), and Boltz-2 agreement of failure modes of other design methods versus LigandForge, our results compare favorably and exceed computational benchmarks of all published binder and peptide generation methods studied herein.

For context, absolute PPI binding affinity prediction remains an unsolved problem across all scoring methods. On published benchmarks: Rosetta REF2015 (InterfaceAnalyzer dG_separated) achieves r = 0.48–0.51 for absolute ΔG ranking on PDBbind-scale datasets, requiring 1–5 minutes per complex for scoring alone and up to 20 minutes with FastRelax.^36^ FoldX AnalyseComplex achieves r = 0.34 on the AB-Bind absolute affinity benchmark, improving to r = 0.37–0.42 on SKEMPI single-point ΔΔG, with 5–10 minutes per complex after RepairPDB.^37^ PRODIGY, a contact-counting method requiring no relaxation, reports r = 0.73 on a curated 79-complex benchmark^7^ but achieves r = 0.34 (95% CI: 0.27–0.40) on the heterogeneous PPB-Affinity dataset we use here. MM/GBSA and MM/PBSA methods achieve r = 0.52–0.75 on small peptide-protein sets but require hours of molecular dynamics simulation per complex.

DeltaForge’s r = 0.83 (95% CI: 0.74–0.89) on curated peptide complexes, at milliseconds per complex, compares favorably within this landscape and is uniquely suited to the high-throughput scoring regime that LigandForge’s generation speed demands. A limitation is that Rosetta and FoldX require institutional licenses (University of Washington and CRG, respectively), preventing us from running them on identical test sets; the comparisons above reference published benchmarks on comparable but not identical datasets.

We note that large-scale efforts are advancing structure-based binder design and affinity estimation. AlphaProteo^25^ demonstrated picomolar-affinity protein binders (50–140 AA) with experimental validation across seven targets; PXDesign^23^ reported 17–82% nanomolar hit rates on six targets via coordinate diffusion; and the Isomorphic Labs Drug Design Engine (IsoDDE) reported small-molecule binding affinity predictions exceeding physics-based methods (r = 0.85 on FEP+ benchmarks). These systems represent important advances but operate in fundamentally different paradigms: all structure-space methods (AlphaProteo, PXDesign, RFDiffusion) generate 3D coordinates followed by inverse folding, requiring iterative structure prediction at inference; IsoDDE is a prediction engine that does not generate novel molecules. The throughput gap is substantial: RFDiffusion requires ∼55 seconds per binder design; RFdiffusion3 is ∼10× faster (∼5–6 sec/design) but still ∼4,000× slower than LigandForge (1.4 ms/design on B200).

AlphaProteo and PXDesign have not published inference throughput. LigandForge occupies a distinct niche: structure-free peptide sequence generation (20–70 AA) conditioned on pocket geometry with thermodynamic supervision compiled into model weights, producing hundreds of candidates per second without structure prediction. These approaches are complementary: LigandForge-generated peptide sequences can be structurally validated by Boltz-2, AlphaFold-family models, or IsoDDE, and the per-residue thermodynamic supervision during training addresses a gap that post-hoc scoring — however accurate — cannot fill.

### Next steps

The immediate next step is experimental validation. Binding studies using SPR and BLI are planned on *a priori*tized subset of high-confidence candidates across multiple target classes to calibrate computational predictions against measured affinities. These experiments will establish the positive predictive value of the dual-metric (iPSAE + ΔG) classification and enable refinement of DeltaForge’s scoring models with experimental ground truth.

Several computational directions will extend this work. First, scaling: LigandForge’s throughput enables systematic exploration across thousands of targets simultaneously, moving from single-target campaigns to portfolio-wide peptide design across entire receptor families. Second, structure-aware training: LigandForge v7.x incorporates explicit 3D coordinate prediction and torsion flow integration during training (while maintaining structure-free inference), enabling the model to learn from backbone geometry without requiring structure prediction at runtime. Early results from LigandForge v7.x indicate that structure prediction can be integrated into the generation pass at ∼20 ms per peptide, with predicted ΔG and Kd consistent with independent Boltz-2 + DeltaForge validation (manuscript in preparation). Third, integration with delivery vehicle optimization: coupling LigandForge peptide design with LNP and other delivery carrier formulation parameters enables co-optimization of binding affinity and pharmacokinetic properties. Fourth, multi-objective optimization: extending the thermodynamic supervision framework to jointly optimize binding affinity, selectivity, and stability within the diffusion process.

The throughput advantage demonstrated here, combined with structure-free generation that compiles thermodynamic supervision into model weights rather than recovering it through iterative search, suggests that amortized, single-pass design may replace iterative optimization as the default paradigm for peptide generation. The 490,691 peptides generated across 150 targets in this work represent, to our knowledge, the largest campaign of computationally designed and structurally validated *de novo* peptides reported to date.

## Methods

### LigandForge Architecture

LigandForge is a generative model based on discrete masked diffusion in amino acid token space.^18, 19^ The production model (v6.5) contains 16.8M trainable parameters (reported as ∼23.7M including optimizer states) and operates on a vocabulary of 24 tokens (20 standard amino acids plus PAD, START, END, and MASK). The architecture comprises three major components:

Pocket Encoder (4.16M parameters, 24.8%). A 4-layer transformer (d=384, 8 attention heads, FFN width=512) encodes the receptor binding pocket from a 48-dimensional per-residue feature vector capturing one-hot amino acid identity, secondary structure, pLDDT, B-factor, formal charge, hydrophobicity, normalized coordinates, and distance to pocket center. The output is a 384-dimensional latent representation per pocket residue with learned position embeddings.

Sequence Denoiser (10.92M parameters, 65.0%). A 6-layer transformer with CROSS-ATTENTION to the pocket latent at every layer (d=384, 8 attention heads, FFN width=768). During training, amino acid tokens are corrupted by a cosine noise schedule, replacing true amino acids with MASK tokens. The model learns to reconstruct the original tokens conditioned on the pocket representation and a continuous time-step embedding. Betancourt & Thirumalai (1999) B-Matrix 20×20 residue pair potentials provide pair-level conditioning. At inference, generation proceeds from a fully masked sequence in a single forward pass (one-shot generation), with nucleus sampling (top-p=0.92) for diversity.

Energy Predictor (2.49M parameters, 14.8%). Comprises four sub-modules: InteractionPredictor (peptide→receptor cross-attention producing contact probability and contact energy matrices), IntraMotifPredictor (within-peptide stability assessment), SecondaryStructurePredictor (Chou-Fasman propensity enforcement with Conv1d layers), and global energy aggregation heads predicting binding ΔG, stability, confidence, and composition quality.

#### Multiscale Thermodynamic Supervision

The key architectural innovation is simultaneous supervision at multiple scales through six loss components: sequence diffusion (cross-entropy), binding energy (MSE + sign agreement on predicted ΔG), interaction contacts (position-weighted MSE on peptide→receptor contact maps with per-residue energy decomposition), binding position enforcement (penalizing zero contacts at known binding residues), intra-peptide stability (MSE on within-peptide contacts and stability score), and composition quality (penalizing unnatural amino acid distributions). The model learns the direct mapping from pocket geometry to energetically favorable amino acids DURING diffusion, not post-hoc. This dense, physically grounded training signal at every scale is why 16.8M parameters achieve competitive results without requiring structure prediction at inference.

#### Training Data

LigandForge was trained on 95,062 bioinspired peptide–receptor complexes derived from a proprietary library of ∼360,000 designed peptides spanning 997 transmembrane protein targets with experimentally characterized protein–protein interaction data. Receptor structures were drawn from ∼3,500 X-ray crystallographic entries in the Protein Data Bank, ensuring ground-truth structural context for all training supervision.

Peptide–receptor complexes were generated using an earlier version of DeltaForge combined with a proprietary Interaction Clipper and *in silico* mutagenesis pipeline.^34^ The Interaction Clipper extracts binding domain fragments from known protein–protein interfaces and maps them onto target receptor pockets; the mutagenesis tool then optimizes these fragments for binding affinity and stability. Each complex was annotated with per-residue and per-atom thermodynamic supervision targets: hydrogen bond energies and geometries, van der Waals contact distances, salt bridge positions, per-residue interaction energies, and global binding ΔG. After quality filtering (removing complexes with steric clashes, broken backbone geometry, or degenerate energy values), 95,062 training examples remained.

The preprocessed dataset (45 GB memory-mapped NumPy arrays) encodes 48-dimensional per-residue receptor pocket features (v6.5 production model), peptide sequences (mean length 37 residues, range 5–150), and multi-scale energy supervision targets at the atomic, residue, sequence, and structural levels. The training corpus spans 2,797 unique X-ray crystallographic PDB structures across 997 receptor targets (230,173 train / 5,772 validation / 6,335 test entries, stratified by receptor).

#### Generalization and Training Overlap

The training corpus contains 2,797 crystallographic PDB structures. At inference, LigandForge uses a structure resolution hierarchy that selects the best available receptor structure per target — crystallographic where available, otherwise AlphaFold or Boltz-2 multimer predictions. Some benchmark targets therefore use the same PDB entry present in training (e.g., TNF-α/1TNF, HER2/1N8Z, KIT/2E9W), while others use structures absent from the training set (e.g., PD-L1/3BIK, VEGF-A/2VPF, IL-7Rα/3UP1). Critically, even when the receptor structure is shared, the training peptides are bioinspired designs from the Interaction Clipper — not the generated peptides being evaluated. Moreover, the generated sequences show no artifacts of training data memorization: across 37,310 unique CD8A peptides, exhaustive pairwise comparison found a maximum similarity of 39.1% (mean 4.5%, zero pairs ≥50%), and across 150,000 peptides spanning five benchmark targets, 100% were unique with per-target maximum similarity of 41.7%. Performance is comparable across targets with and without training-set receptor overlap (Table 4a), consistent with generalization of binding physics rather than memorization of specific receptor–peptide pairs.

#### Optional Mask Channels

LigandForge supports optional conditioning masks: binding mask (residues expected to contact the receptor), stability mask (residues contributing to intramolecular stability), linker mask (flexible connecting segments), and specialized masks for glycosylation sites, ion coordination, and water bridge positions.

### DeltaForge Scoring Engine

DeltaForge is implemented in Rust with Python bindings via PyO3, using Rayon for parallelism. For each protein-protein or protein-peptide complex, DeltaForge computes 17 structural features: hydrogen bond count, salt bridge count, hydrophobic contacts, six PRODIGY-style contact classification counts (CC, CP, CA, PP, PA, AA), pi-pi stacking count, cation-pi interaction count, water bridge count, shape complementarity score, conformational entropy cost, and interface size. Binding free energy (ΔG) is predicted using size-bin-specific linear models fitted by Huber regression on the PPB-Affinity benchmark dataset. Kd is derived from ΔG at inference via the thermodynamic relation ΔG = RT·ln(Kd) at 298K for interpretability; Kd is never used as a training objective because the exponential ΔG-to-Kd transformation compresses the gradient signal and distorts the loss landscape.

### PPB-Affinity Benchmark Validation

DeltaForge was validated against the PPB-Affinity dataset,^8^ which aggregates experimental binding affinity data from five curated databases: SKEMPI v2.0,^9^ PDBbind v2020,^10^ SAbDab,^11^ ATLAS,^12^ and Affinity Benchmark v5.5.^13^ After filtering for crystal structures, assigning chains, and removing entries with zero detected contacts, we obtained 4,347 complexes from 2,848 unique PDB structures. Cross-validation was performed within each size bin using 5-fold splits stratified by source database.

### Boltz-2 Validation Pipeline

Generated peptides were validated by folding in complex with target receptor structures using Boltz-2.^16^ We compute iPSAE (interface predicted Structural Alignment Error) from the PAE matrix as the mean alignment confidence across all residue pairs where one residue is in the peptide and the other is in the receptor, normalized to.^0, 1^ Where pre-folded receptor structures were available, Boltz-2 was run in template mode: the receptor structure was provided as a template with constrained backbone, reducing folding time by 10–40-fold.

### Model Versions and Training Details

The results reported in this paper were generated using LigandForge v6.5, the production model deployed for inference. LigandForge v6.5 is a 16.8M-parameter model (reported as ∼23.7M including optimizer states) with a 4-layer pocket encoder (4.16M params; 48-dim input, 384-dim latent), a 6-layer sequence decoder with 8-head cross-attention to the pocket at every layer (10.92M params), and an energy predictor (2.49M params) with six loss components. Training used 95,062 preprocessed protein-peptide complexes with DeltaForge-computed energy decompositions as supervision targets. All analyses were performed using Python 3.11, PyTorch 2.x, Rust 1.75+, and Boltz-2 v0.4.

## Software and Data Availability

LigandForge, DeltaForge, and H-SWAE are proprietary software of Ligandal, Inc. Access to the LigandForge peptide design platform is available as a public beta at https://ligandai.com. The DeltaForge scoring engine is available as a web API for academic evaluation upon request. The PPB-Affinity benchmark dataset is publicly available.^8^

## Supplementary Tables

**Table S1:**
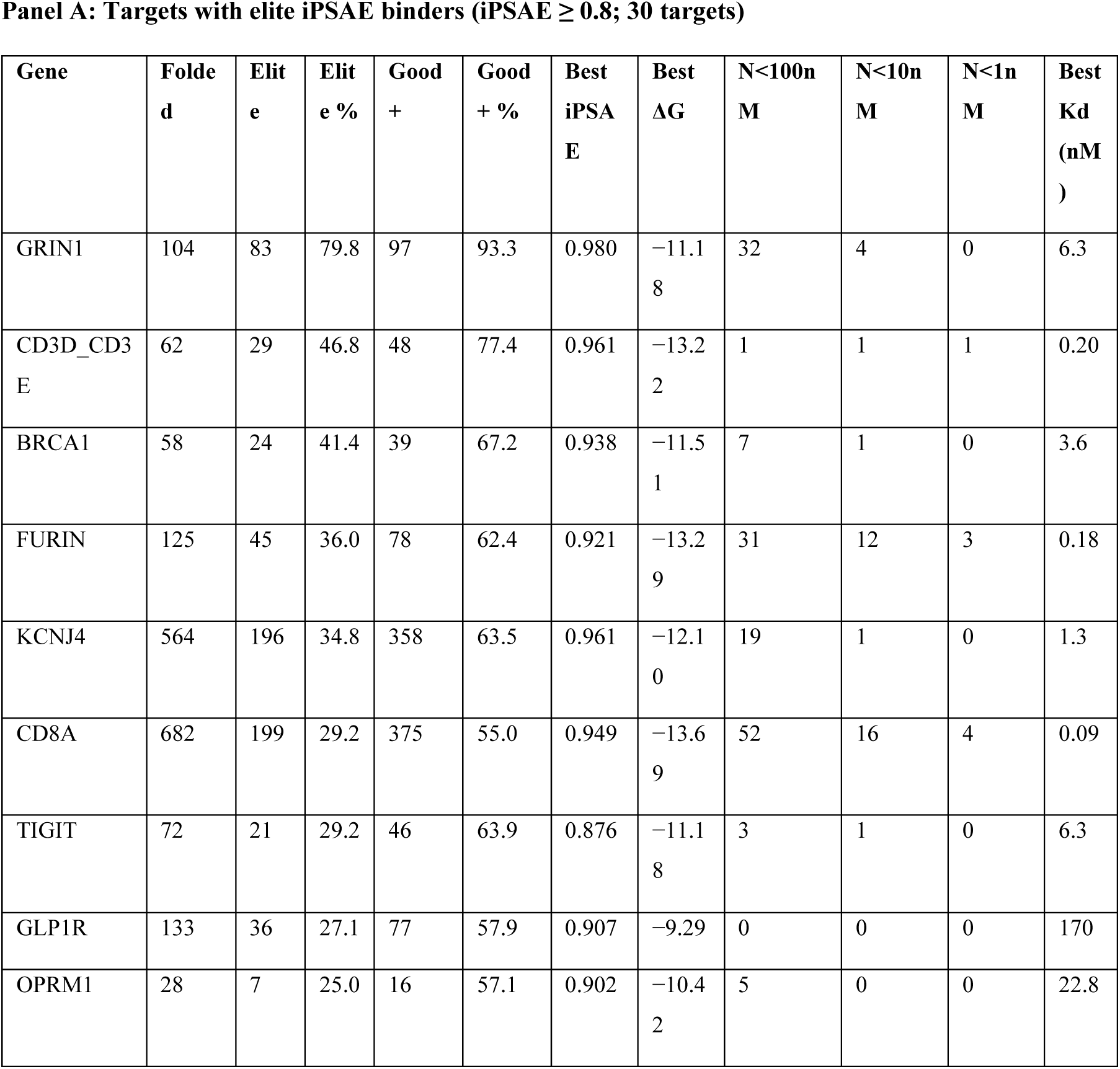
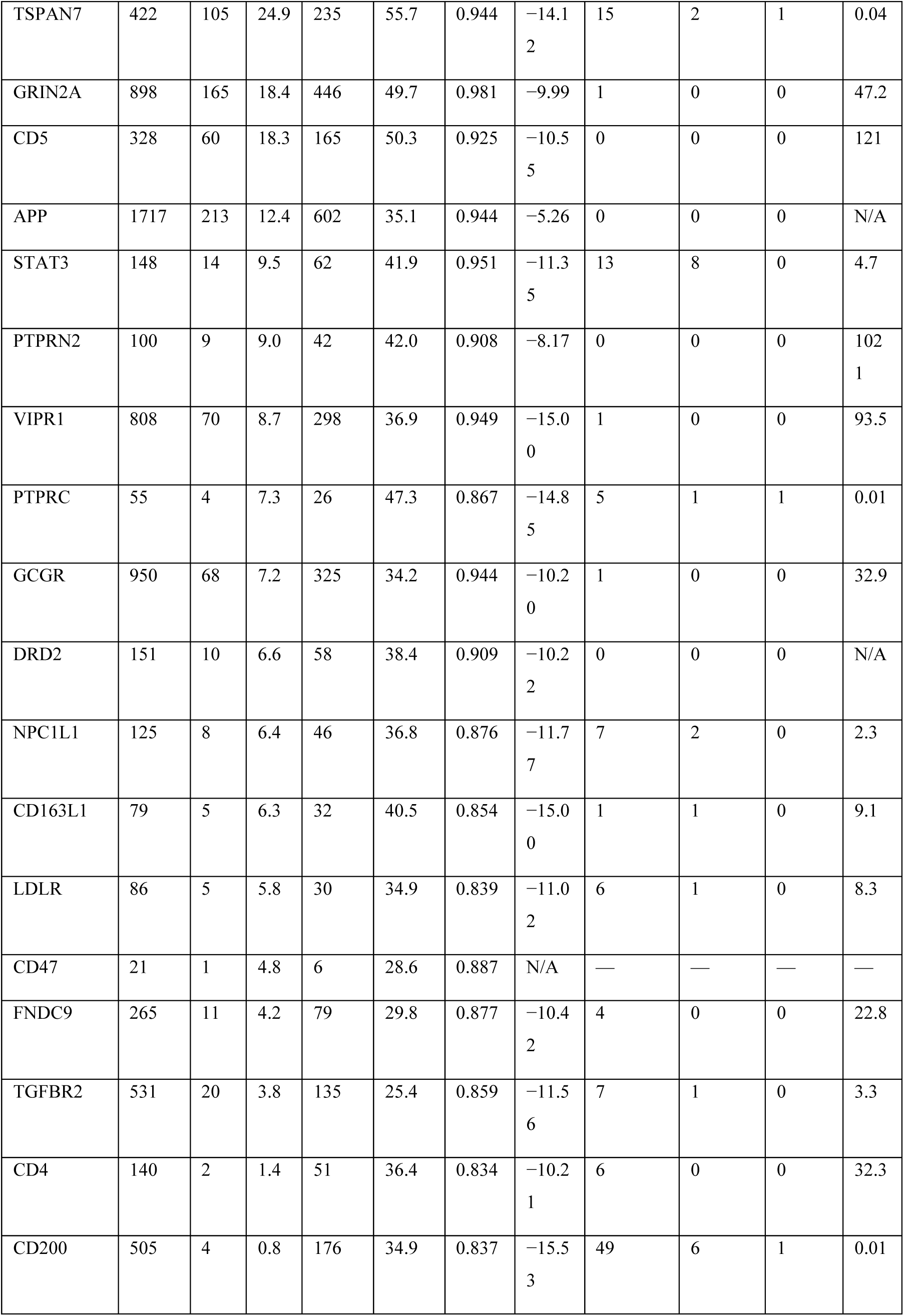
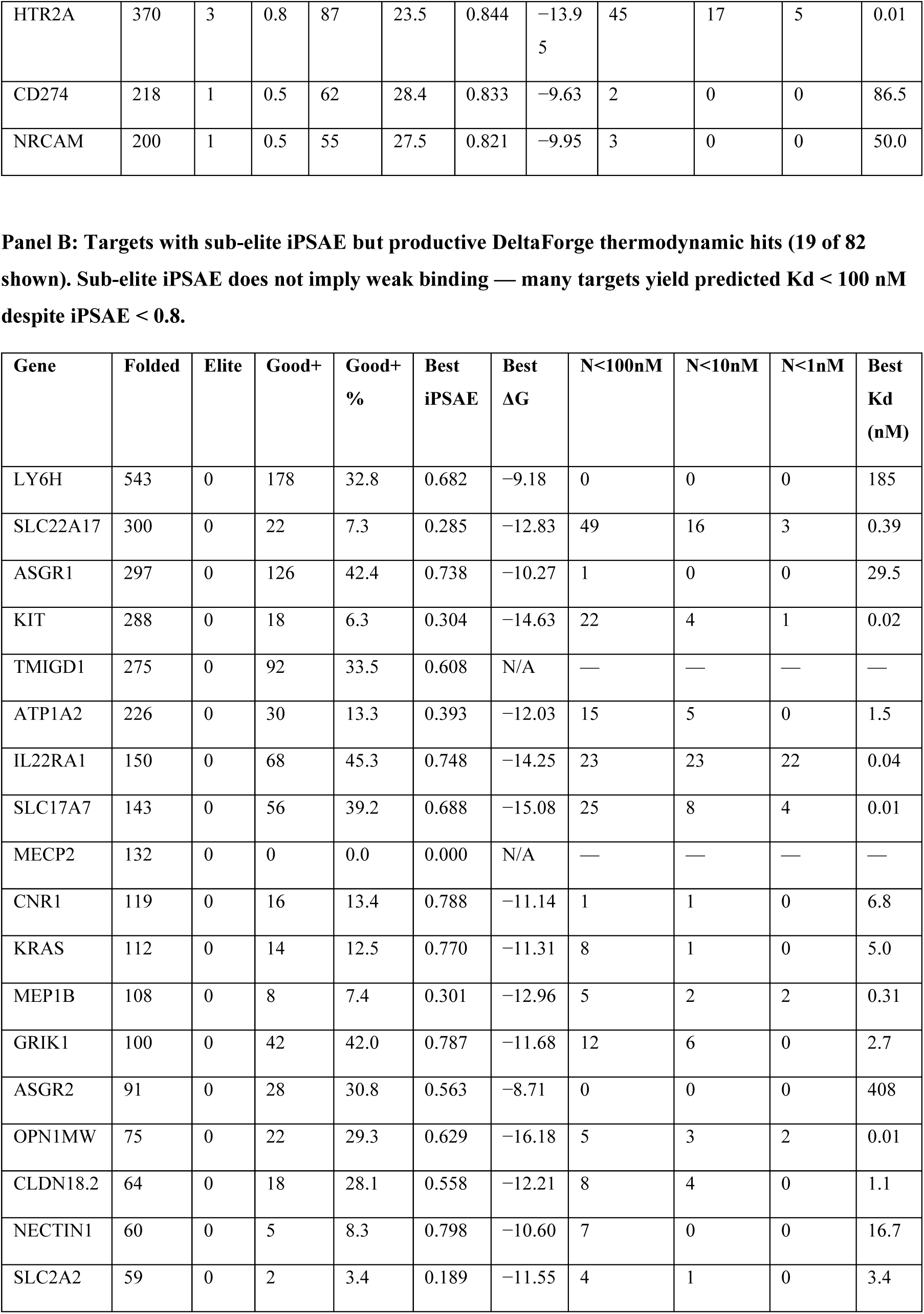
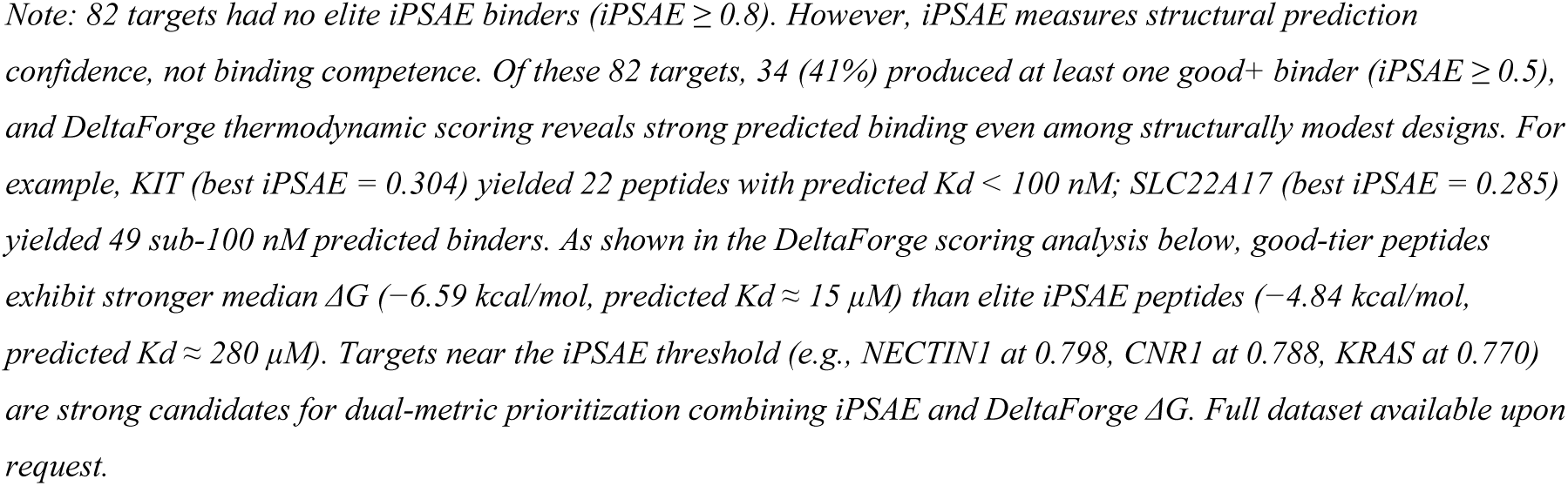
Cumulative Campaign. Complete target-level summary across the LigandForge campaign, sorted by elite iPSAE binder percentage (descending), then by total folded (descending). Elite defined as iPSAE >= 0.8; Good+ defined as iPSAE >= 0.5. DeltaForge ΔG predictions from Boltz-2-folded structures are shown alongside iPSAE metrics; Kd bin columns indicate the number of scored peptides with predicted dissociation constant below each threshold. N/A or — indicates no DeltaForge scoring was completed for that target. Note: 20 targets have a best ΔG at the DeltaForge prediction floor (−15.0 kcal/mol); sub-1 nM counts for these targets should be interpreted with caution.

**Table S2:** MD Relaxation of Representative Complexes. Molecular dynamics relaxation results for the nine representative structures shown in Figure 8. Each structure was relaxed using OpenMM steepest descent minimization with AMBER14 force field and Cα harmonic restraints (k = 10 kcal/mol/Å², 100 steps). Campaign best Kd values represent the strongest DeltaForge-predicted binder across all scored peptides for each target (not limited to the displayed structure). N<100 nM and N<1 nM columns show the total count of predicted sub-threshold binders per target.

| Target | iPSAE | Clashes<br>(before) | Clashes<br>(after) | Ca<br>RMSD<br>($\text{\AA}$) | Converged | $N < 100$<br>nM | $N < 1$ nM | Best $K_d$<br>(nM) |
| --- | --- | --- | --- | --- | --- | --- | --- | --- |
| CD8AB | 0.862 | 15 | 2 | 0.119 | Yes | 127 | 73 | $\leq 0.01^*$ |
| CD3D–<br>CD3E | 0.974 | 11 | 4 | 0.099 | Yes | 1 | 1 | 0.20 |
| GRIN1 | 0.980 | 8 | 8 | 0.000 | No | 32 | 0 | 6.3 |
| HTR2A | 0.844 | 19 | 2 | 0.088 | Yes | 45 | 5 | $\leq 0.01^*$ |
| FURIN | 0.921 | 21 | 4 | 0.088 | Yes | 31 | 3 | 0.18 |
| KCNJ4 | 0.961 | 19 | 0 | 0.096 | Yes | 19 | 0 | 1.3 |
| TIGIT | 0.876 | 2 | 1 | 0.088 | Yes | 3 | 0 | 6.3 |
| CD5 | 0.925 | 12 | 4 | 0.116 | Yes | 0 | 0 | 121 |
| CD3E–<br>CD3G | 0.970 | 17 | 17 | 0.001 | No | — | — | — |

\* Best $K_d$ at DeltaForge prediction floor ( $\Delta G = -15.0$ kcal/mol); exact value should be interpreted with caution. — indicates DeltaForge scoring not available for this target.

## Competing Interests

A.W. is the founder, Chief Executive Officer, and a member of the board of directors of Ligandal, Inc. LigandForge, DeltaForge, LigandAI, and Predictive Interactomics are proprietary technologies of Ligandal, Inc. Patent applications covering the LigandForge discrete diffusion architecture, DeltaForge thermodynamic scoring engine, and related methods have been filed by Ligandal, Inc.

*This manuscript describes computational methods and predictions. All binding affinity estimates are computational and require experimental validation. LigandForge and DeltaForge are trademarks of Ligandal, Inc*.

## Footnotes

† Throughout this paper, † marks values at the DeltaForge prediction floor (−15.0 kcal/mol per chain). The floor reflects the limit of the scoring model’s calibrated range, not a claim of ultra-high affinity; reported Kd values derived from floor-capped ΔG should be treated as upper bounds (i.e., the true Kd is at most this strong, and may be weaker if the ΔG is an overestimate).

## References

1. Muttenthaler, M. et al. Trends in peptide drug discovery. Nat. Rev. Drug Discov. 20, 309–325 (2021).

2. Lau, J.L. & Dunn, M.K. Therapeutic peptides: Historical perspectives. Bioorg. Med. Chem. 26, 2700–2707 (2018).

3. Pacesa, M. et al. One-shot design of functional protein binders with BindCraft. Nature (2025). doi:10.1038/s41586-025-09429-6.

4. Stark, H. et al. BoltzGen: toward universal binder design. bioRxiv (2025). doi:10.1101/2025.11.20.689494.

5. Watson, J.L. et al. De novo design of protein structure and function with RFdiffusion. Nature 620, 1089–1100 (2023).

6. Abdin, O. & Kim, P.M. Target sequence-conditioned design of peptide binders. Nat. Biotechnol. (2025). doi:10.1038/s41587-025-02761-2.

7. Xue, L.C. et al. PRODIGY: predicting binding affinity of protein-protein complexes. Bioinformatics 32, 3676–3678 (2016).

8. Renaud, N. et al. PPB-Affinity: A benchmark for protein-protein binding affinity prediction. Sci. Data (2024).

9. Jankauskaite, J. et al. SKEMPI 2.0. Bioinformatics 35, 462–469 (2019).

10. Wang, R., et al. The PDBbind database. J. Med. Chem. 48, 4111–4119 (2005).

11. Dunbar, J. et al. SAbDab: the structural antibody database. Nucleic Acids Res. 42, D1140–D1146 (2014).

12. Berman, H.M. et al. ATLAS. Database (2022).

13. Vreven, T. et al. Docking Benchmark Version 5 and Affinity Benchmark Version 2. J. Mol. Biol. 427, 3031–3041 (2015).

14. Uhlen, M. et al. Tissue-based map of the human proteome. Science 347, 1260419 (2015).

16. Wohlwend, J., et al. Boltz-2: Biomolecular structure prediction for every molecule. Preprint (2025).

17. Kastritis, P.L. & Bonvin, A.M. Are scoring functions ready to predict interactomes? J. Proteome Res. 9, 2216–2225 (2010).

18. Austin, J. et al. Structured denoising diffusion models in discrete state-spaces. NeurIPS (2021).

19. Sahoo, S.S., et al. Simple and effective masked diffusion language models. NeurIPS (2024).

20. Kolouri, S., et al. Sliced Wasserstein auto-encoders. ICLR (2019).

21. Bryant, P. et al. Structure prediction of protein-peptide complexes. Nat. Commun. (2024).

22. Gainza, P. et al. Accurate de novo design of high-affinity protein-binding macrocycles. Nat. Chem. Biol. (2025).

23. Guo, Z., et al. PXDesign: Fast, modular, and accurate de novo design of protein binders. bioRxiv (2025).

24. Friedman, D. & Dieng, A.B. The Vendi Score. Trans. Mach. Learn. Res. (2023).

25. Zambaldi, V. et al. De novo design of high-affinity protein binders with AlphaProteo. Nature (2025).

26. Ariax Bio. FreeBindCraft: open-source BindCraft fork replacing PyRosetta with OpenMM. GitHub: cytokineking/FreeBindCraft (2025).

27. Nori, J. et al. BindEnergyCraft: replacing ipTM with pTMEnergy for protein binder design. arXiv:2505.21241 (2025). ICML 2025.

28. Krishna, R. et al. RFdiffusion2: generalized biomolecular design with all-atom structure prediction. Nat. Methods (2025).

29. Watson, J.L., et al. RFdiffusion3: universal generative model for biomolecular design. Preprint (2025).

30. Bryant, P., et al. Blind de novo design of dual cyclic peptide agonists targeting GCGR and GLP1R. bioRxiv (2025). doi:10.1101/2025.06.06.658268.

31. Krumm, B.E. et al. De novo design of miniprotein agonists and antagonists targeting G protein-coupled receptors. Science (2025). doi:10.1126/science.adx0080.

32. Schoenmakers, P. et al. Assessment of generative de novo peptide design methods for GPCRs. bioRxiv (2026). doi:10.64898/2026.02.26.708415.

33. Dunbrack, R.L. Rēs ipSAE loquunt: What’s wrong with AlphaFold’s ipTM score and how to fix it. bioRxiv (2025). doi:10.1101/2025.02.10.637595.

34. Watson, A., et al. Peptide antidotes to SARS-CoV-2 (COVID-19). bioRxiv (2020). doi:10.1101/2020.08.06.238915.

35. Overath, R., et al. Predicting experimental success in de novo binder design: a meta-analysis of 3,766 experimentally characterised binders. bioRxiv (2025). doi:10.1101/2025.08.14.670059.

36. Alford, R.F. et al. The Rosetta All-Atom Energy Function for Macromolecular Modeling and Design. J. Chem. Theory Comput. 13, 3031–3048 (2017).

37. Schymkowitz, J. et al. The FoldX web server: an online force field. Nucleic Acids Res. 33, W382–W388 (2005).

38. Alamdari, S., et al. Protein generation with evolutionary diffusion: sequence is all you need. bioRxiv (2023). doi:10.1101/2023.09.11.556673.

39. Li, Z., et al. Full-atom peptide design with geometric latent diffusion. ICML (2024).

40. Roney, J.P., Ou, C. & Ovchinnikov, S. Protein diffusion models as statistical potentials. bioRxiv (2025). doi:10.64898/2025.12.09.693073.

